# Iterative inhibition of commissural growth cone exploration, not post-crossing barrier, ensures forward midline navigation through SlitC-PlxnA1 signaling

**DOI:** 10.1101/2020.04.21.051359

**Authors:** Hugo Ducuing, Thibault Gardette, Aurora Pignata, Karine Kindbeiter, Muriel Bozon, Olivier Thoumine, Céline Delloye-Bourgeois, Servane Tauszig-Delamasure, Valérie Castellani

## Abstract

Sensitization to Slits and Semaphorin (Sema)3B floor plate repellents after midline crossing is thought to be the mechanism expelling commissural axons contralaterally and preventing their back-turning. We studied the role of Slit-C terminal fragment sharing with Sema3B the Plexin (Plxn) A1 receptor, newly implicated in midline guidance. We generated a knock-in mouse strain baring PlxnA1Y1815F mutation altering SlitC but not Sema3B responses and observed recrossing phenotypes. Using fluorescent reporters, we found that Slits and Sema3B form clusters decorating an unexpectedly complex mesh of ramified FP glia basal processes spanning the entire navigation path. Time-lapse analyzes revealed that impaired SlitC sensitivity destabilized axon trajectories by inducing high levels of growth cone exploration from the floor plate entry, increasing risk of aberrant decisions. Thus, FP crossing is unlikely driven by post-crossing sensitization to SlitC. Rather, SlitC limits growth cone plasticity and exploration through reiterated contacts, continuously imposing a straight and forward-directed trajectory.

## Introduction

Extensive efforts have been put in the past decades into the identification of guidance cues and the receptors mediating their effects on navigating axons (Raper and Mason, 2010). Combinatorial expression of guidance cues at choice points along the path as well as versatile perception of these cues by growth cones were shown to largely contribute to coding the diversity of guidance decisions underlying the wiring of neuronal circuits (Raper and Mason, 2010; Stoeckli et al, 2019; Ducuing et al, 2019). In the recent years, a number of reports brought the evidence that guidance receptors have multiple binding partners, assembling into macro-complexes at the growth cone membrane under the influence of intrinsic or extrinsic triggers, and interacting in addition with more than a single ligand (Seiradake et al, 2016). This scheme significantly complexifies our understanding of axon guidance processes, making the characterization of functional outcomes downstream of each receptor module a central challenge. Noticeably too, the guidance sources controlling navigating axons appear to be much more diversified than anticipated. For instance, cell migration streams were found to deliver guidance molecules to navigating axons at meeting points (Lopéz-Bendito et al, 2011). Bipolar progenitors were reported to contribute to commissural axon navigation by transporting Netrin on their basal processes, which then accumulates on the pial surface and is thought to serve as guidance support for axons pathing in its vicinity (Dominici et al, 2017; Varadarajan et al, 2017). These studies also highlight a lack of knowledge concerning the modes of action of the guidance cues. Indeed, how they are presented to the growing axons and how their patterns of distribution are regulated remain poorly addressed questions, leaving open whether the ligand topography takes a determinant part in the generation of specific guidance decisions.

The navigation of commissural axons across the spinal cord midline provides a very suited model to address these questions. Commissural axons navigate the midline ventrally in the floor plate (FP), guided by a wide range of cues. We reported in previous work that PlexinA1 (PlxnA1) is a receptor shared by different FP repulsive cues. When associated with Neuropilin2 (Nrp2), it mediates the response to Semaphorin3B (Sema3B) (Nawabi et al, 2010), and while unbound to Nrp2, it acts as a receptor for a second type of ligands, the Slit1-3-C (Delloye-Bourgeois et al, 2015). These C terminal fragments are generated from the processing of integral Slit1-3 proteins together with Slit1-3-N terminal fragments, whose effects are mediated by Roundabout (Robo) receptors (Evans and Bashaw, 2010). It turns out that at least three signaling pathways are thought to generate repulsive forces in the FP. They are considered indispensable to push the axons in the contralateral side and also to set a midline barrier preventing any turning back (Ducuing et al, 2019).

Genetic deletions of ligands or receptors to abrogate individual signaling were shown to alter the ability of commissural growth cones to exit the FP and interpreted as if repulsive forces were no longer sufficient to drive the growth forwards and prevent midline recrossing. Moreover, these phenotypic analyses also pointed out differences in guidance errors, suggesting that individual signaling pathways might regulate distinct aspects of the commissural navigation. Invalidation of Robo-SlitN or PlxnA1-Nrp2-Sema3B signaling were both reported to impair FP crossing by inducing stalling of commissural axons within the FP. An additional aberrant behavior - axons turning back towards the ipsilateral side - was found in mice lacking either PlxnA1 or all Slit ligands. This indirectly suggested that the PlxnA1-SlitC signaling might specifically set the midline barrier that prevents commissural axons from crossing back (Ducuing et al, 2019, Chedotal, 2019).

In addition, various experiments suggested that the perception of repulsive forces is set at an appropriate timing, in order to synchronize ligands activity with commissural axon progression (Neuhaus-Follini and Bashaw, 2015; Pignata et al, 2016). Experimental manipulations leading to premature sensitivity were shown to induce commissural axons to stall at the FP entry or to undergo an ipsilateral turn (Chen et al, 2008; Nawabi et al, 2010; Ducuing et al, 2019). We recently reported that responsiveness to Slits and Sema3B is sequentially acquired through successive sorting of Nrp2, PlxnA1, Robo1, and Robo2 receptors at the growth cone membrane during the FP navigation (Pignata et al, 2019). This study however still leaves open several puzzling questions: What is the nature of the guidance forces generated by these signaling during the progression of commissural axons within the FP? How are the instructions from the different guidance cues present in the FP translated into directional information for axon growth? In particular, whether physical and/or chemical substrates build a midline barrier once the axons have crossed is still fully hypothetic. It is unclear as well whether commissural axons move away under the influence of ligands gradients. Moreover, so far, rare attention has been given to the topology of the FP territory in which commissural axons path, although increasing attention is given to physical properties exerted by environmental substrates during the formation of neuronal projections (Kerstein et al, 2015). FP glial cells are thought to have a typical progenitor-like bipolar morphology, with a dorsal apical anchor at the lumen of the central canal and a basal process laying ventrally on the basal lamina, even though some studies suggested these cells have much more complex morphology (Campbell et Peterson, 1993). The axon path is dorsally bordered by the soma and ventrally by the basal lamina (Figure 1A). Despite seminal studies performed with electronic microscopy, which described close contacts between the axons and glial cells (Yaginuma et al. 1991), little is known about the exact morphology of the basal processes and the putative physical constraints that axons might have to face during their FP navigation.

**Figure 1:**
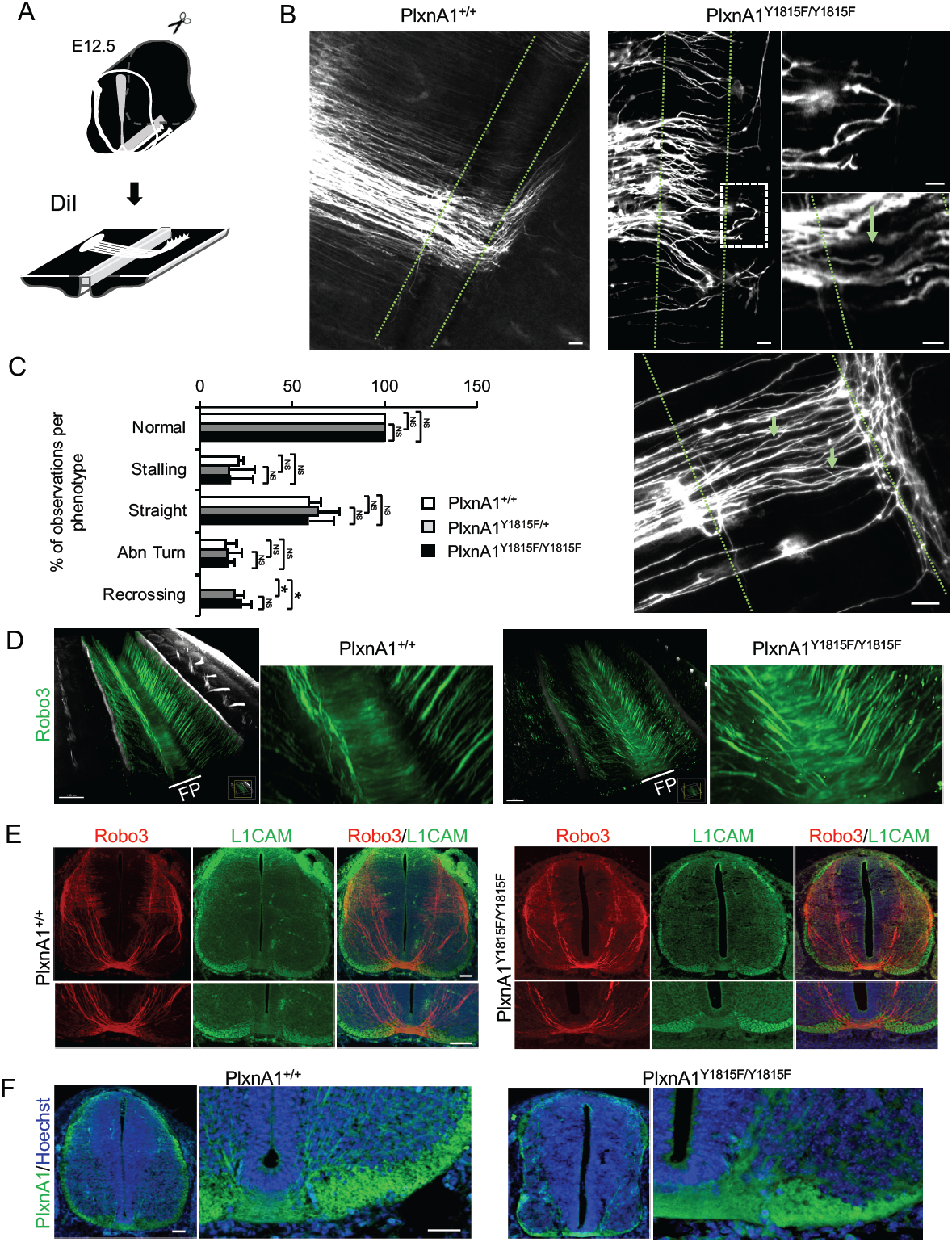
Y1815F mutation in PlxnA1 induces commissural axon recrossing and disorganized trajectories. (A) Schematic drawing of open-book preparations for DiI tracing of commissural axon trajectories. (B) (left panel) Microphotographs illustrating commissural tracts extending straight towards the FP, crossing the FP, and turning rostrally at the FP exit in PlxnA1^+/+^ embryos. (right panel) Microphotographs and magnifications illustrating the phenotypes observed in the PlxnA1^Y1815F/Y1815F^ embryos, with axons turning back during the navigation, and misdirected trajectories within the FP (indicated by green arrows). The FP is delimited by dashed green lines. (C) Quantitative analysis of commissural axon phenotypes showing that the Y1815F mutation induces recrossing, while it does not affect other aspects of the navigation such as the stalling (PlxnA1^+/+^, N = 3 embryos; PlxnA1^+/Y1815F^, N = 5 embryos; PlxnA1^Y1815F/Y1815F^, N = 4 embryos). Data are shown as the mean ± s.e.m, Student test has been applied, *: P-value p < 0.05. (D) Light sheet imaging of the spinal commissural tracts in PlxnA1^+/+^ and PlxnA1^Y1815F/Y1815F^ embryos at E12.5, immunostained with anti-Robo3 antibody. Note the disorganized aspect of commissural axons crossing the FP. (E) Immunofluorescent labeling of Robo3 and L1CAM in transverse cryosections from E12.5 embryos illustrating the general patterns of pre-crossing commissural axons (revealed by Robo3 marker) and post-crossing axons (revealed by L1CAM marker) in PlxnA1^+/+^ and PlxnA1^Y1815F/Y1815F^ spinal cords. (F) PlxnA1 immunolabeling on E12.5 transverse sections of PlxnA1^+/+^ and PlxnA1^Y1815F/Y1815F^ embryos at E12.5 showing that the general pattern of the receptor is preserved in the mutated context. Scale bar: 10µm in (B), 50µm in (D, E, F).

In the present study, we focused on the SlitC-PlxnA1 signaling and investigated its functional outcome during commissural axon navigation. Through combinations of *in vivo* analysis, live imaging and super resolution microscopy, we show that growth cone trajectories are actively kept straight over the FP navigation, through reiterated contacts with SlitC spots presented by ramifications of FP glia basal process that mark out the entire FP navigation path. PlxnA1 mutation altering perception of SlitC alleviates these constrains, resulting in highly dynamic growth cones that actively explore their local environment, which compromises efficient and straight navigation across the FP. This reveals that guidance forces preventing midline recrossing are not generated by post-crossing sensitization to repellents, but through continuous limitation of growth cone exploration to restrict the navigation direction forward.

## Results

### PlxnA1^Y1815F^ mutation in mice recapitulates PlxnA1^-/-^ recrossing but not stalling phenotype

We identified in previous work a tyrosine residue of PlxnA1 cytoplasmic tail, whose phosphorylation was specifically triggered upon SlitC exposure (Delloye-Bourgeois et al, 2015). Expression of a mutant receptor form for this tyrosine, PlxnA1^Y1815F^, in commissural neurons significantly reduced their growth-cone collapse response to SlitC, while their sensitivity to Sema3B was preserved (Delloye-Bourgeois et al, 2015). The presence of re-crossing phenotype in embryos lacking PlxnA1 or all Slit1-3 but not Robo1/2 or Sema3B led us to speculate that PlxnA1-SlitC signaling might provide the specific force that prevents commissural axons from turning back in the FP. To test this model, we generated a PlxnA1^Y1815F^ mouse strain. Homozygous individuals were viable, fertile, having no obvious morphological alterations (Supplemental Figure 1A-D). PlxnA1 expression in the spinal cord was further validated in western blot on lysates of spinal cords isolated from PlxnA1^+/+^, PlxnA1^Y1815F/+,^ PlxnA1^Y1815F/Y1815F^ and PlxnA1^-/-^ embryos (Supplemental Figure 1E). To examine commissural axon trajectories, we traced the commissural tracts with fast 1,1′-dilinoleyl-3,3,3′,3′-tetramethylindocarbocyanine, 4-chlorobenzenesulfonate (DiI) crystals inserted in the dorsal domain of open-book mounted embryonic spinal cords isolated from E12.5 embryos having +/+, Y1815F/+, and Y1815F/Y1815F genotypes. Axon trajectories were classified according to previous work (Delloye-Bourgeois et al, 2015). Interestingly, while no difference of stalling was observed between PlxnA1^+/+^ and PlxnA1^Y1815F/Y1815F^ embryos, the recrossing cases were exclusively associated with loss of one or two copies of the PlxnA1 gene, reaching the proportion reported in PlxnA1^-/-^ embryos (Figure 1A-C). Moreover, in the Y1815F context, axon trajectories appeared disorganized, with axons baring a tortuous aspect, even in cases where they succeeded to cross the FP and reached the post-crossing compartment (Figure 1B).

To further examine how the Y1815F mutation affects commissural axon navigation, E12.5 embryos were immunolabeled with an antibody against Robo3 commissural marker, cleared, and imaged with light sheet microscopy. We observed in 3D analysis a clear disorganization of the Robo3^+^ commissural tracts in PlxnA1^Y1815F/ Y1815F^ embryos compared with wild-type littermates, with axons losing their straight and parallel trajectory when navigating the FP (3 embryos for each genotype, Figure 1D).

Thus, the ability of commissural axons to maintain a straight trajectory towards the FP exit requires a functional PlxnA1-SlitC signaling. In contrast, the lack of stalling phenotype in PlxnA1^Y1815F/Y1815F^ embryos indicates that the guidance forces resulting from the unaffected PlxnA1-Nrp2-Sema3B signaling remain sufficient for pushing commissural axons towards the FP exit.

### PlxnA1^Y1815F^ mutation does not obviously affect pre-crossing commissural navigation

To examine the pattern of commissural axon projections prior to the FP crossing, transverse sections of E12.5 embryos were immunostained with antibodies directed against known pre-crossing and post-crossing markers of commissural axons, such as Robo3 and L1CAM, respectively. Analysis of commissural axon patterns in confocal images revealed no obvious defects of coursing towards the FP, indicating that PlxnA1 Y1815 residue is dispensable for the pre-crossing navigation (Figure 1E). This is consistent with our previous finding that PlxnA1 is not delivered at the commissural growth cone surface prior to the FP entry, and is thus not expected to have prominent role at the pre-crossing stage (Pignata et al, 2019). The expression pattern of PlxnA1 was also not obviously different between WT and PlxnA1^Y1815F/Y1815F^ embryos, being detected in both cases at highest levels in crossing and post-crossing axons (Figure 1F). This indicated that the Y1815 mutation does not prevent neither the synthesis of PlxnA1 receptor nor its trafficking to the axon compartment.

### PlxnA1^Y1815F^ expressing axons fail to maintain forward directed growth when navigating the FP

To gain insights into how the alteration of SlitC-PlxnA1 signaling impacts on the behavior of navigating axons, we first thought to re-introduce WT and mutated versions of PlxnA1 in E12.5 PlxnA1^-/-^ murine embryos by *ex vivo* electroporation of their neural tube. We constructed a pHluo tagged version of PlxnA1^Y1815F^ receptor by fusing the pHluo to the N-terminal part of the coding sequence to take advantage of this pH sensitive GFP allowing reporting the presence of the receptor at the cell membrane (Pignata et al, 2019). The pHluo-PlxnA1^Y1815F^ and pHluo-PlxnA1^WT^ constructs were co-electroporated along with a mbTomato construct (Figure 2A). We took advantage of our previous work in which we compared the outcome of different plasmid concentrations on the FP navigation to select a concentration of 2µg/µl, allowing sufficient receptor expression level and navigation across the FP (Pignata et al, 2019). The pH dependency of the pHluo constructs was verified as in Pignata et al, (2019). Right after electroporation, the spinal cords were dissected out and open-books were incubated for two days and imaged at fixed time-point with a spinning disk confocal microscope to examine the aspect of commissural axon trajectories in the FP. We analyzed individual axon trajectories spanning the FP by counting the number of deviations they presented and classifying them from O (straight and slightly and regularly curved with no deviation angle), 1, 2 and up to 2 deviation angles. Interestingly, we observed that while pHluo-PlxnA1^WT+^growth-cones exhibited a 73,4% of straight growth trajectories, this ratio dropped to 35,6% for commissural axons that had sorted PlxnA1^Y1815F^ at their surface, that displayed much more irregular curly aspects, with trajectories having significantly increased number of curvature (Figure 2B, C). Video-time lapse recording on these electroporated mouse open-books turned out to be highly challenging, mainly due to high and fast toxicity resulting from repeated laser exposure. We thus turned to the chicken embryo as an alternative model for video-microscopy of commissural axon navigation in a context of impaired SlitC signaling.

**Figure 2:**
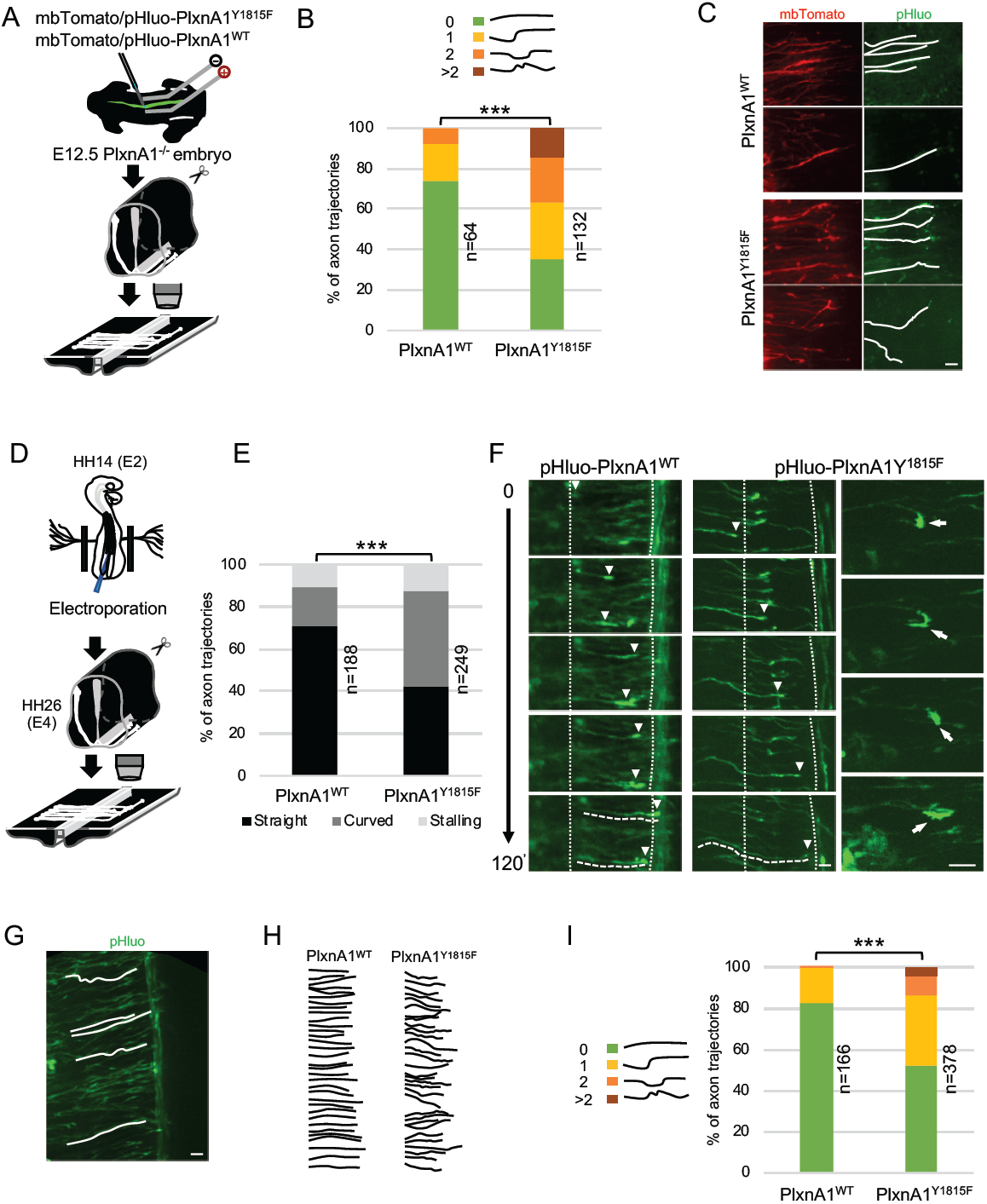
Commissural neurons expressing PlxnA1^Y1815F^ at their surface fail to maintain straight growth during FP navigation. (A) Schematic drawing of the paradigm of electroporation of pHluo-PlxnA1 receptor forms in PlxnA1^-/-^ mouse embryonic spinal cord and open-book preparations for live imaging. (B) Histogram depicting the analysis of axon trajectories, classified by counting curvatures according to the indicated criteria, showing that straight trajectories are less frequent when commissural neurons express PlxnA1^Y1815F^ than PlxnA1^WT^ (PlxnA1^WT^, N = 10 embryos, 64 growth cones; PlxnA1^Y1815F^, N = 7 embryos, 132 growth cones). Chi^2^ test has been applied, ***: P-value p < 0.001. (C) Microphotographs illustrating navigating commissural growth cones expressing the mutated pHluo-receptor at their surface, reported by the green pHluo fluorescent signal. The red signal depicts the mbTomato, co-electroporated with the pHluo-receptor construct. Note that some axons have a curly aspect. (D) pHluo-receptor constructs were co-electroporated with the mbTomato in the chicken neural tube and spinal cords were mounted in open-books for time-lapse imaging. (E) Histogram quantifying axon trajectories reconstructed from time-lapse sequences of pHluo^+^ growth cones navigating the FP, showing increased proportion of wavy axon shapes in the PlxnA1^Y1815F^ condition, compared with PlxnA1^WT^ one (PlxnA1^WT^, N = 3 embryos, 188 growth cones; PlxnA1^Y1815F^, N = 3 embryos, 249 growth cones). Chi^2^ test has been applied, ***: P-value p < 0.001. (F) Snapshots of the movies illustrating the commissural growth patterns (white triangles) and an illustration of a PlxnA1^Y1815F+^ growth cone turning back. The FP is delimited by dashed white lines. (G, H) Microphotograph illustrating traces of commissural axon trajectories. Example of traces patterns from several snapshots. (I) Histogram depicting the quantification and showing increased curvatures of axon shapes in the PlxnA1^Y1815F^ condition compared with PlxnA1^WT^ one (PlxnA1^WT^, N = 3 embryos, 166 growth cones; PlxnA1^Y1815F^, N = 3 embryos, 378 growth cones). Chi^2^ test has been applied, ***: P-value p < 0.001. Scale bars: 10µm in (C, F, G).

We thus electroporated pHluo-PlxnA1^WT^ and pHluo-PlxnA1^Y1815F^ in the neural tube of the chicken embryo and performed time lapse analysis of the commissural axon navigation as in Pignata et al (2019). We first plotted the trajectories of individual axons at successive time points, classifying them as straight or curved. We also took advantage of live imaging to assess growth cone stalling. We found in the PlxnA1^WT^ condition a majority of axons having a straight trajectory (79,7%), whereas in contrast PlxnA1^Y1815F +^ axons exhibited a more tortuous aspect in 64,5% of the cases (Figure 2D-F). The rate of growth cone stalling was comparable in the WT and Y1815F conditions, which was also consistent with the lack of increased stalling phenotype in PlxnA1^Y1815/Y1815^ embryos, compared to wild-type littermates. Finally, expressing PlxnA1^Y1815F^ in chicken commissural neurons also resulted in pictures of growth cones turning back (Figure 2F). As done in our experiments with mouse open-books, we also analyzed commissural axon trajectories to quantify their degree of deviation. Consistent with our previous observations, we found the percentage of straight trajectories significantly decreased from 82.7% in the PlxnA1^WT^ condition to 52.6% in the PlxnA1^Y1815^F one, with correlated increase in curved shapes (Figure 2G-I).

Altogether these experiments confirmed that PlxnA1^Y1815F^ is unable to ensure the forward growth direction that is normally taken by commissural axons to cross the FP and validated the use of the chicken embryo as a model for further investigations.

### The temporal and spatial pattern of PlxnA1 membrane insertion during commissural axon navigation is impacted by the Y1815 mutation

Next, we investigated whether the Y1815 mutation alters the SlitC-PlxnA1 signaling by modifying the dynamics of PlxnA1 in commissural growth cones in living conditions. First, we analyzed the pattern of pHluo receptor introduced in PlxnA1^-/-^ mouse embryos. As expected from our immunohistochemical labeling of PlxnA1 in PlxnA1^Y1815F^ transverse sections, pHluo-PlxnA1^Y1815F^ was trafficked to the axon and the growth cones. However, the cell surface patterns of PlxnA1^Y1815F^ and PlxnA1^WT^ strikingly differed in axons navigating the FP. While PlxnA1^WT^ was mostly restricted to the growth cone, PlxnA1^Y1815F^ occupied a much larger membrane domain, overflowing in the adjacent axon shaft compartment (Figure 3A-C). The difference was statistically significant, as shown by measures of the area of pHluo signal in both PlxnA1^Y1815F^ and PlxnA1^WT^ axons (Figure 3B). Thus, the Y1815 mutation alters the cell surface pattern of PlxnA1 receptor in navigating commissural axons.

**Figure 3:**
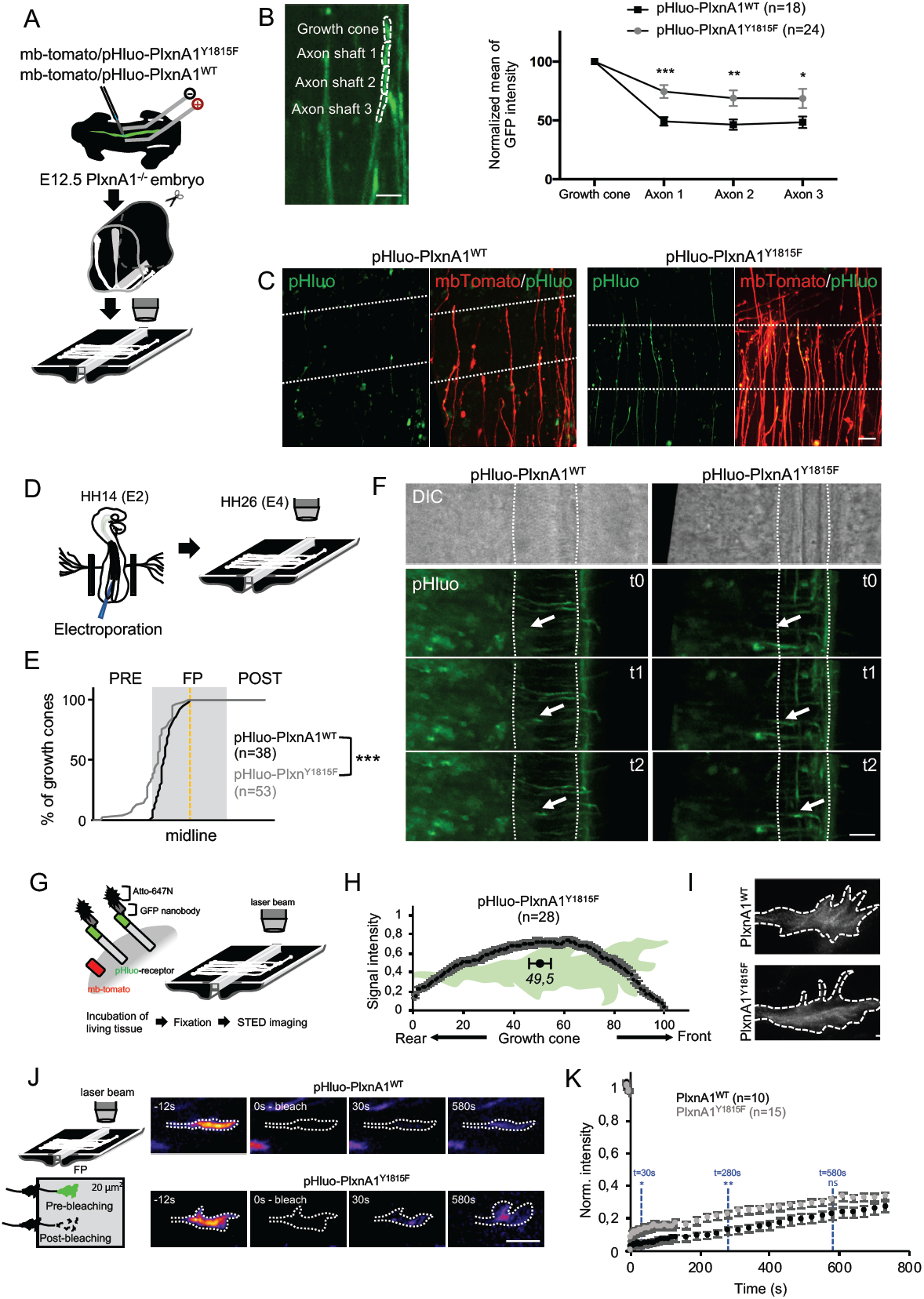
the Y1815F mutation alters the spatio-temporal pattern of cell surface PlxnA1 distribution and membrane mobility at the growth cone. (A) Schematic drawing of the paradigm of electroporation of the mouse embryonic spinal cord. (B) Method and quantification of the pHluo signal in commissural axons navigating the floor plate (FP), showing an expansion of the expression domain in the PlxnA1^Y1815F^ condition, compared with the PlxnA1^WT^ one (PlxnA1^WT^, N = 6 electroporated PlxnA1^-/-^ embryos, 24 growth cones; PlxnA1^Y1815F^, N = 4 PlxnA1^-/-^ electroporated embryos, 18 growth cones). Data are shown as the mean ± s.e.m, Student test has been applied, *: P-value p < 0.05, **: P-value p < 0.01, ***: p < 0.001. (C) Microphotographs of commissural axons navigating the FP in living open-books, illustrating the difference of PlxnA1 cell surface distribution between the WT and Y1815F condition. The FP is delimited by dashed white lines. (D) Schematic drawing of the experimental paradigm of expression of PlxnA1 in the chicken spinal cord. (E) Cumulative fractions of growth cones that sort PlxnA1 to their surface, reported by the pHluo fluorescence (pHluo-PlxnA1^WT^, N = 2 electroporated embryos, 38 growth cones; pHluo-PlxnA1^Y1815F^, 4 electroporated embryos, 53 growth cones). In the PlxnA1^Y1815F^ condition, 100% of the growth cone population has sorted the receptor when reaching the midline, but for a substantial amount of the growth cones, the sorting had already occurred prior to the FP entry. KS test has been applied, ***: P-value p < 0.001. (F) Microphotographs of live imaging movies illustrating the sorting of PlxnA1 receptor at the surface of commissural growth cones navigating the FP. The phase contrast image depicts the position of the FP in the open-books (delimited by dashed white lines). The green signal corresponds to the pHluo fluorescence, reporting the presence of PlxnA1 at the membrane. In the WT condition, PlxnA1 is sorted at the membrane from the FP entry to the midline. (G) Schematic representation of the paradigm of STED imaging in open-books. (H) Quantification of the center of mass of the signal, showing no polarity towards the rear or the front of the growth cones (N= 28 imaged growth cones from 5 electroporated embryos). Data are shown as the mean ± s.e.m. (I) STED microscopy images of pHluo-PlxnA1^Y1815F^ cell surface distribution in commissural growth cones navigating the FP. The signal distributes over the entire growth cone surface. (J) Schematic representation of the paradigm of FRAP experiments in open-books. Representative color-coded images from video time-lapse sequences illustrating photobleaching and fluorescence recovery of fluorescent growth cones navigating the FP. (K) Graphs of fluorescence recovery for pHluo-PlxnA1^WT^ and pHluo-PlxnA1^Y1815F^ (PlxnA1^WT^, N = 2 embryos, 10 growth cones; PlxnA1^Y1815F^, N = 2 embryos, 15 growth cones). Data are shown as the mean ± s.e.m, Student test has been applied at t=30s, t=280s, and t=580s, ns: non-significant, *: P-value p < 0.05, **: P-value p < 0.01. Scale bars: 5µm in (I), 10µm in (B, C, J), 50µm in (F).

Second, we recently reported that PlxnA1 is specifically delivered at the growth cone surface when commissural axons navigate the first part of the FP, a timing that differs from that of Robo1, which is sorted later on, in the second FP half (Pignata et al, 2019). To examine whether the Y1815 mutation affects the temporal sequence of PlxnA1 membrane insertion, the pHluo-PlxnA1^Y1815F^ and pHluo-PlxnA1^WT^ constructs, along with the mbTomato construct, were electroporated in the neural tube of chicken embryos. Receptor dynamics were studied in video-microscopy of commissural axons navigating the FP in open-books. We analyzed the temporal pattern of membrane insertion by pointing the position of growth cones switching on the pHluo fluorescence. Interestingly, we found with this cartography analysis that in both PlxnA1^WT^ and PlxnA1^Y1815F^ conditions, nearly 100% of the growth cones navigating the FP had sorted the receptor before midline crossing, indicating that the temporal pattern of membrane insertion within the FP was unaffected by the Y1815 mutation (Figure 3D-F, movies S1-S2). Nevertheless, we also observed that a significant proportion of PlxnA1^Y1815F^- growth cones, that could navigate the FP, had sorted the receptor prior to the FP entry (Figure 3E, movies S3-S4). This suggested that the mutated receptor might be sorted precociously, although it appeared not to arrest the growth cones at the FP entry.

Third, in recent work using this experimental paradigm, we observed using STED microscopy that the membrane pool of PlxnA1^WT^ concentrates at the front of the growth cone. Thus, we thought to examine how PlxnA1^Y1815F^ distributes at the surface of the commissural growth cone. Living open-books electroporated with mbTomato/pHluo-PlxnA1^Y1815F^ constructs were incubated with Atto-647N-conjugated GFP nanobodies to label cell surface pHluo. Open-books were fixed, and plxnA1^Y1815F^ was imaged in growth cones navigating the FP. Notably, in sharp contrast with our observations of PlxnA1^WT^, we observed that PlxnA1^Y1815F^ distributed from the front to the rear. This apparent absence of polarity was confirmed by measurement of the signal intensity along the rear-front axis (Figure 3G-I).

Thus, overall, these analyses showed that PlxnA1^Y1815F^ spatial distribution in commissural axons navigating the FP is altered, which reveals that the mutation might interfere with SlitC-specific traffic and signaling mechanisms.

### Y1815F results in increased membrane mobility of PlxnA1 receptor in navigating commissural growth cones

To gain further insights into the biological process impaired by theY1815F mutation, we conducted FRAP experiments to compare exocytosis and membrane motility of WT and mutated PlxnA1 in commissural growth cones navigating the FP. The pHluo-receptor fluorescence was bleached to 80-90% at time zero, in a 15 to 20μm^2^ area, which allows covering the entire growth cone surface. Then, the fluorescence recovery was recorded for 20 min (Figure 3J, K, movies S5-S8). Strikingly in the PlxnA1^Y1815F^ condition, the fluorescence recovery was significantly higher over the first time points, than in the PlxnA1^WT^ one, and the difference established from this early step remains constant over time. The temporality of the recovery pattern difference indicates that the PlxnA1^Y1815F^ has a faster membrane diffusion in the membrane than wild-type receptor. The stability of the difference over time suggests in addition that exocytosis events might be unaffected by the mutation. Thus, Y1815F mutation results in relapse of PlxnA1 motility at the cell membrane which, either upstream or downstream of SlitC binding, affects the ability of the receptor to mediate SlitC signal.

### FP glia cells elaborate complex ramified basal processes staking the axon path

Next, we investigated how SlitC maintains a straight growth of commissural axons. To gain insights, we first thought to get a precise analysis of the topology of the axon navigation path in the FP and to characterize the spatial distribution of the different ligands (Figure 4A). We first proceeded to an immunostaining of transverse sections of E4 chick embryos with an antibody recognizing the FP glial cell marker Ben. The staining revealed an unexpected dense network of processes in the basal compartment corresponding to the axon path, in particular with BEN+ structures aligned in the left-right axis (Figure 4B). We next performed a sparse electroporation of chick embryos with a plasmid encoding a membrane-bound mbTomato to observe individual cells. We set-up a position of the electrodes on both sides of E2 embryos, allowing to specifically target FP glial cells. Spinal cords were dissected two days later and imaged in open-books by confocal microscopy (Figure 4C). We observed that the glial cell elaborates a morphologically complex basal process (Figure 4D, E). 3D reconstruction with IMARIS software after deconvolution treatment of the images revealed a “squid-like” structure, with a basal process subdivided into several pillars, with typically several anchored to the ventral pole (Figure 4F). The pillars appeared much larger in the left-right axis than in the rostro-caudal one, as suggested by previous data from electronic microscopy (Okabe et al, 2004; Figure 4E-F). To examine the commissural axon path, we stained E4 spinal cord open-books with NgCAM commissural marker, and FP cells with the nuclear DAPI staining. In the rostro-caudal axis, we observed a “ladder rungs”-type NgCAM staining with rows of nuclei regularly interspaced, that delineated rostro-caudal interspaced streams through which the axons navigate. In the dorso-ventral axis, axons were constrained by the nuclei in the basal compartment (Figure 4G-H). To examine the dorso-ventral organization of commissural axons, we measured the intensity of NgCAM staining in dorsal (basal A) and ventral (basal B) equal sub-compartments (Figure 4I-J). We found that NgCAM staining was slightly enriched in the more ventral compartment, suggesting more axons are taking a ventral route (Figure 4J). Next, we electroporated a mbTomato plasmid containing a Math1 promoter, which specifically drives the expression in dorsal commissural neurons (Helms, 1998) and proceeded to immuno-staining of electroporated spinal cords with an anti-BEN antibody. This allowed us to visualize more precisely that commissural growth cones are individually intercalated between glial cell processes (Figure 4K). We finally co-electroporated Math1-mbTomato plasmid with a plasmid encoding GFP under the control of HoxA1 promoter, which specifically drives the expression in FP glial cells (Li and Lufkin, 2000), (Figure 4L-M). From confocal images of Math1-mbTomato fluorescent signal we reconstructed the morphology of commissural growth cones. We observed them infiltrating the space between pillars, having an oblong shape, flattened in the left-right dimension and elongated in the dorso-ventral one (Figure 4N). This analysis revealed that the FP glia builds highly complex mesh of basal processes whose stereotypic spatial organization provides a physical frame for commissural growth cones, likely imposing numerous and repeated contacts all over the FP navigation (Figure 4O, movies S9-S12).

**Figure 4:**
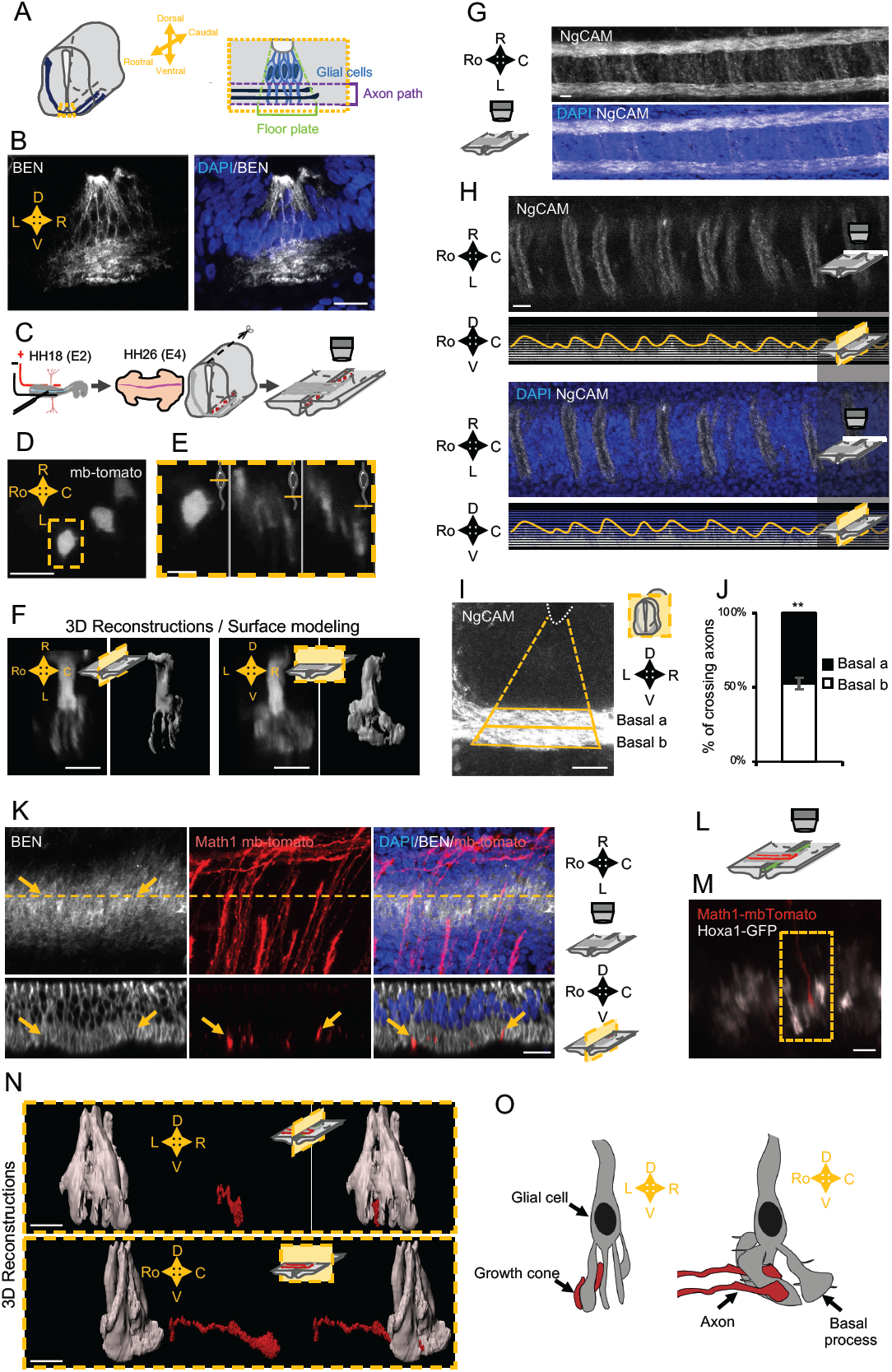
Spatial organization of the commissural axon navigation path. (A) Schematic drawings of a spinal cord at E12.5 when commissural axons cross the floor plate (FP) (left panel) and close-up of the FP (right panel) with glial cells (light blue) and crossing axons (dark blue). (B) Immunostaining of an E4 FP transverse section with FP specific BEN antibody (left panel) and merged with DAPI staining (right panel). (C) *In ovo* FP electroporation procedure. 48h after electroporation, spinal cords are dissected and mounted as open-books for time lapse microscopy. Sparse electroporated mbTomato electroporated cells are visualized in red. (D) Open-book imaging of E4 chick FP with sparse mbTomato electroporation. The dashed square highlights a glial cell. (E) Close-up of the single glial cell observed in (D), at three different positions along the dorso-ventral axis, as shown by the schematic representation on the upper right corner of each image. A single FP glial cell displays multiple basal feet. (F) 3D reconstruction (first and third panels) and surface modeling (second and last panels) of a single FP cell seen in a sagittal section (left panels) or transverse section (right panels). (G, H) Open-book view of E4 chick spinal cord floor-plate showing crossing axons stained with NgCAM at 20x magnification (first and third panels) and reconstruction of the corresponding sagittal view (second and last panels). In the open-book view, the depth shown is at the most dorsal part of the axon path. (I) 80µm transverse section of E4 chick spinal cord FP showing crossing axons stained with NgCAM. The FP basal domain was subdivided in two and NgCAM intensity was measured. (J) NgCAM in both basal subdomains (N= 3 embryos, 2 sections per embryo, 3 images analyzed per section). Data are shown as the mean ± s.d, paired Student test has been applied, **: P-value p < 0.01. (K) 3D reconstruction of axons navigating through the floor plate stained with DAPI (in blue), BEN (in white) and with mbTomato electroporated in axons (in red). The yellow dashed line corresponds to the cut plane resulting in the sagittal optic section shown in the lower panel. Yellow arrows point growth cones intercalated between BEN labeled glial cell processes when they navigate the FP (see also movies of this 3D reconstruction in Supplemental information movies S9-13). (L) Schematic drawings of an open-book with a sparse electroporation of commissural axons and a broad FP electroporation. Two plasmids are used: Math1-mbTomato drives the expression of a membrane anchored mbTomato in commissural neurons, while HoxA1-GFP drives the expression of GFP in FP glial cells. (M) Ventral longitudinal view from an open-book electroporated as described in (L). The yellow dashed square delineates a close-up of a crossing growth cone electroporated with Math1-mbTomato (in red) navigating along the basal feet of glial cells electroporated with Hoxa1-GFP (in white). (N) Surface reconstruction of a growth cone electroporated with Math1-mbTomato (in red) navigating along the basal feet of glial cells electroporated with Hoxa1-GFP (in white) as seen in a sagittal section (upper panel) or transversal section (lower panel). (O) Schematic drawing of a glial cell and two axons crossing through its basal end-feet, in a sagittal view (left panel) or transverse view (right panel). Scale bars: 5µm in (D, E, F, L, M), 20µm in (B, G, H).

### SlitC and SlitN are expressed as segregated clusters in the FP navigation path

Next, we studied the distribution patterns of PlxnA1 ligands in the FP. For Sema3B, we took advantage of a knock-in model of Sema3B-GFP fusion generated in our previous work (Arbeille et al, 2015). Transverse sections were stained with anti-GFP antibodies to amplify the GFP signal, allowing detection of the protein deposited in the basal domain (Figure 5A). Quantitative analysis of the GFP signal along the mediolateral axis of the basal domain and along the dorso-ventral axis of the FP revealed protein clusters distributed evenly in the FP, arranged in columns reflecting their localization along the basal processes. We then concentrated on Slit proteins. Study of Slit fragments distribution pattern is limited by the lack of antibodies allowing their detection and their distinction from the full-length form. Thus, as in Xiao et al (2011) we developed an alternative strategy based on fluorescent reporters, whose distribution should approximate the pattern of endogenous proteins, since their physico-chemical properties might be highly similar. We constructed a plasmid encoding full-length Slit2 (Slit2 FL), fused to two distinct fluorescent proteins: Cerulean at its N-terminal part and Venus at its C-terminal part. Slit2 FL is visualized with both fluorophores overlapping (white signal). Upon cleavage, Slit N is reported by Cerulean fluorescence (here in purple) and Slit C by Venus (here in green) (Figure 5B). This construct was electroporated in the FP of E2 chicken embryos, which were dissected two days later (Figure 5C). Thick transverse embryonic sections were prepared and the FP observed by confocal microscopy. For analysis, the FP domain was subdivided into three compartments along the dorso-ventral axis: (i) the “apical” one, delimited by the central canal and the bottom of glial cells nuclei, (ii) the “basal a” encompassing the dorsal half of the axons path and (iii) the “basal b”, containing the ventral half of the axons path until the basal lamina (Figure 5D). Slit2 dual-tagged construct allowed us to observe that glial cells display a massive white staining in their apical part, reflecting the presence of the uncleaved Slit2 FL and/or overlapping Slit2N and Slit2C fragments. Conversely, the basal compartments, in particular the most ventral one in which commissural axons preferentially path, showed higher degree of separation of the Slit2 fragments, as quantified by a Pearson coefficient evaluation (Figure 5E). Analysis of cerulean (Slit2N) and Venus (Slit2C) fluorescence showed that both Slit2 fragments had a graded pattern, with higher levels in the basal a *versus* basal b compartment. Yet, Slit2C was significantly enriched in the basal b compartment, when compared to Slit2N (Figure 5F). Thus, Slit2C and Slit2N distribution patterns within the FP territory are close, although they present some specificities which could result from distinct structural properties and/or from active mechanisms regulating Slit2 prior to the generation of the fragments. To address this question, we examined whether Slit2C and Slit2N patterns resulting from endogenous Slit2 processing were similar to those observed when the fragments are expressed individually. We thus constructed two plasmids encoding one fragment each in secreted fusion with a GFP (Figure 5G). The patterns of distribution of Slit2N-GFP and Slit2C-GFP were then analyzed as performed previously. Interestingly, whereas Slit2N displayed the same distribution pattern when expressed as an isolated fragment or generated from Slit2 cleavage, Slit2C diffused massively in the axon path when issued from Slit2 cleavage (Figure 6H-J). Furthermore, Slit2C-GFP isolated fragment was more prone to deposition along the FP basal membrane than Slit 2C processing product (Figure 5K). Thus, beyond physicochemical properties, Slit2C and Slit2N patterns likely result from additional mechanisms regionalizing full-length Slit2 protein and its processing within the FP glial cell compartments.

**Figure 5:**
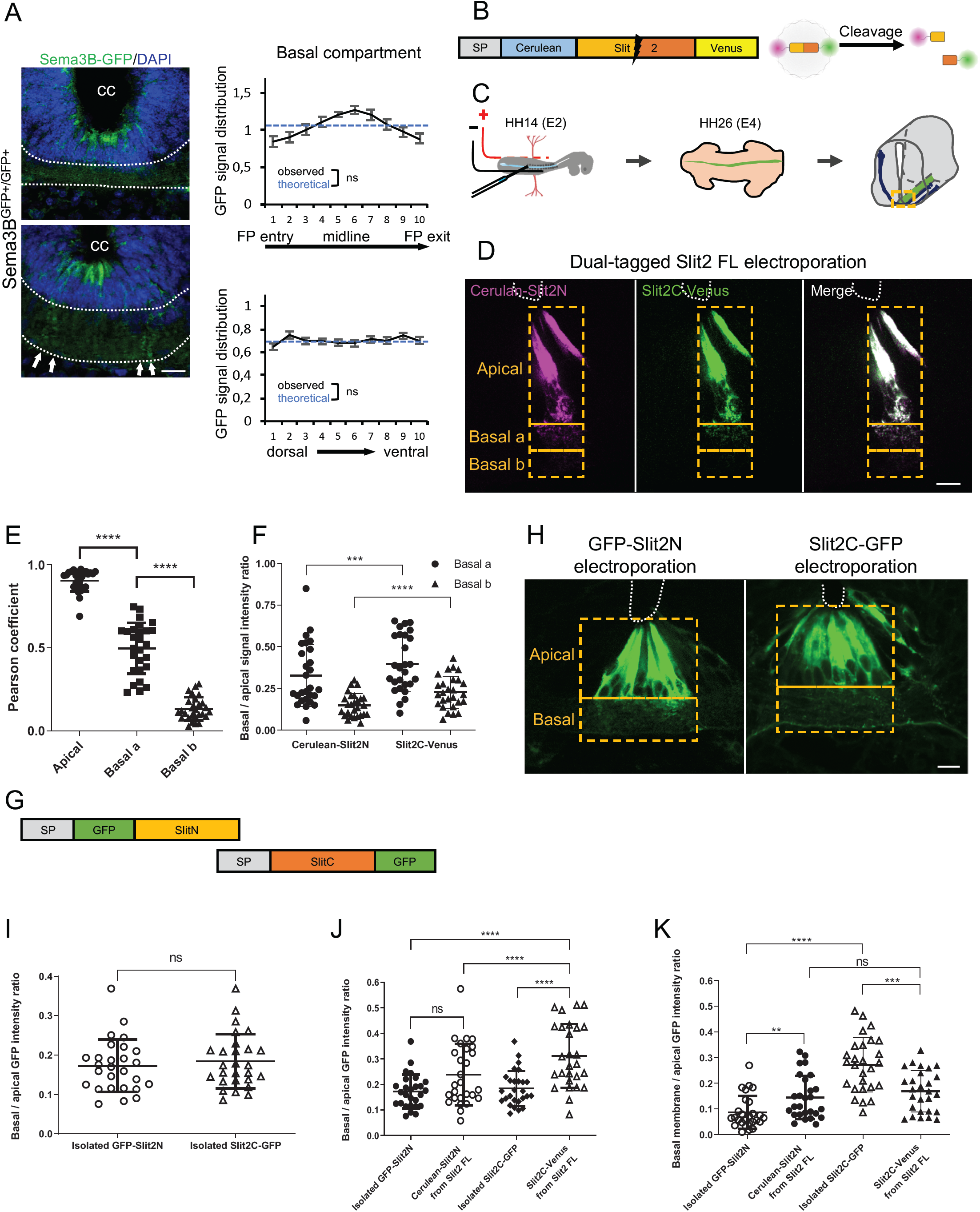
Slit2N and Slit2C decorate the FP glia basal processes and have distinct diffusion properties conditioned by Slit2 FL processing. (A) Transverse section of Sema3B-GFP homozygous E12.5 embryo stained with DAPI. Endogenous GFP reports Sema3B expression and is quantified in the left-right axis of the FP (upper panel) and in the dorso-ventral axis (lower panel). Both distributions do not differ from a homogeneous distribution. The axon path is delineated by white dashed lines and the white arrows indicate the deposition of Sema3B-GFP along the glial cell processes. Data are shown as the mean ± s.e.m, KS test has been applied, n.s. (B) Schematic drawings of dual-tagged Slit2 construct and activity. Dual-tagged Slit2 coding sequence was cloned in pCAGEN vector. Full-length Slit2 (Slit2 FL) is fused to Cerulean at its N-terminal part and Venus at its C-terminal part. Slit2 FL is visualized with both fluorophores overlapping (white signal). Upon cleavage, Slit2N presence is reported by Cerulean fluorescence (here in purple) and Slit2C by Venus (here in green). (C) *In ovo* FP electroporation procedure. 48h after electroporation, embryos were fixed in PFA and sliced on vibratome. (D) 80µm transverse section of E4 chick spinal cord FP electroporated with dual-tagged Slit2. (E) Pearson coefficients quantify the degree of colocalization of Cerulean and Venus in FP electroporated with dual-tagged Slit2, through three compartments: apical, basal a, and basal b, as delimited by the dashed square in (D) (N= 3 embryos, 3 sections per embryo, 3 images analyzed per section). (F) Intensity ratio of the basal compartment over the apical compartment for Cerulean and Venus in FP electroporated with dual-tagged Slit2. (G) Schematic drawings of Slit2 isolated fragments fused to GFP. Both fusion protein coding sequence were cloned under the control of a HoxA1 promoter to drive the expression within the FP glial cells. (H) 80µm transverse sections of E4 chick spinal cord FP electroporated with isolated Slit2 fragments fused to GFP. The apical and basal compartment are delineated with yellow dashed lines. (I) Intensity ratio of the basal compartment over the apical compartment for GFP in FP electroporated with either Slit2N-GFP or Slit2C-GFP (N= 3 embryos, 3 sections per embryo, 3 images analyzed per section). (J) Comparison between the basal/apical intensity ratio of Slit2 isolated fragments compared to the basal/apical intensity ratio of Slit2N and Slit2C fragments generated by the cleavage of dual-tagged Slit2 FL. (K) Comparison between the basal membrane/apical domain intensity ratio of Slit2 isolated fragments compared to the basal membrane/apical domain ratio of Slit2N and Slit2C fragments generated by the cleavage of dual-tagged Slit2 FL. Data are shown as the mean ± s.d. in (E, F, I, J, K), Student test has been applied, ns: non-significant, ***: P-value p < 0.001, ****: p < 0.0001. Scale bars: 10µm in (D, H), 50µm in (A).

**Figure 6:**
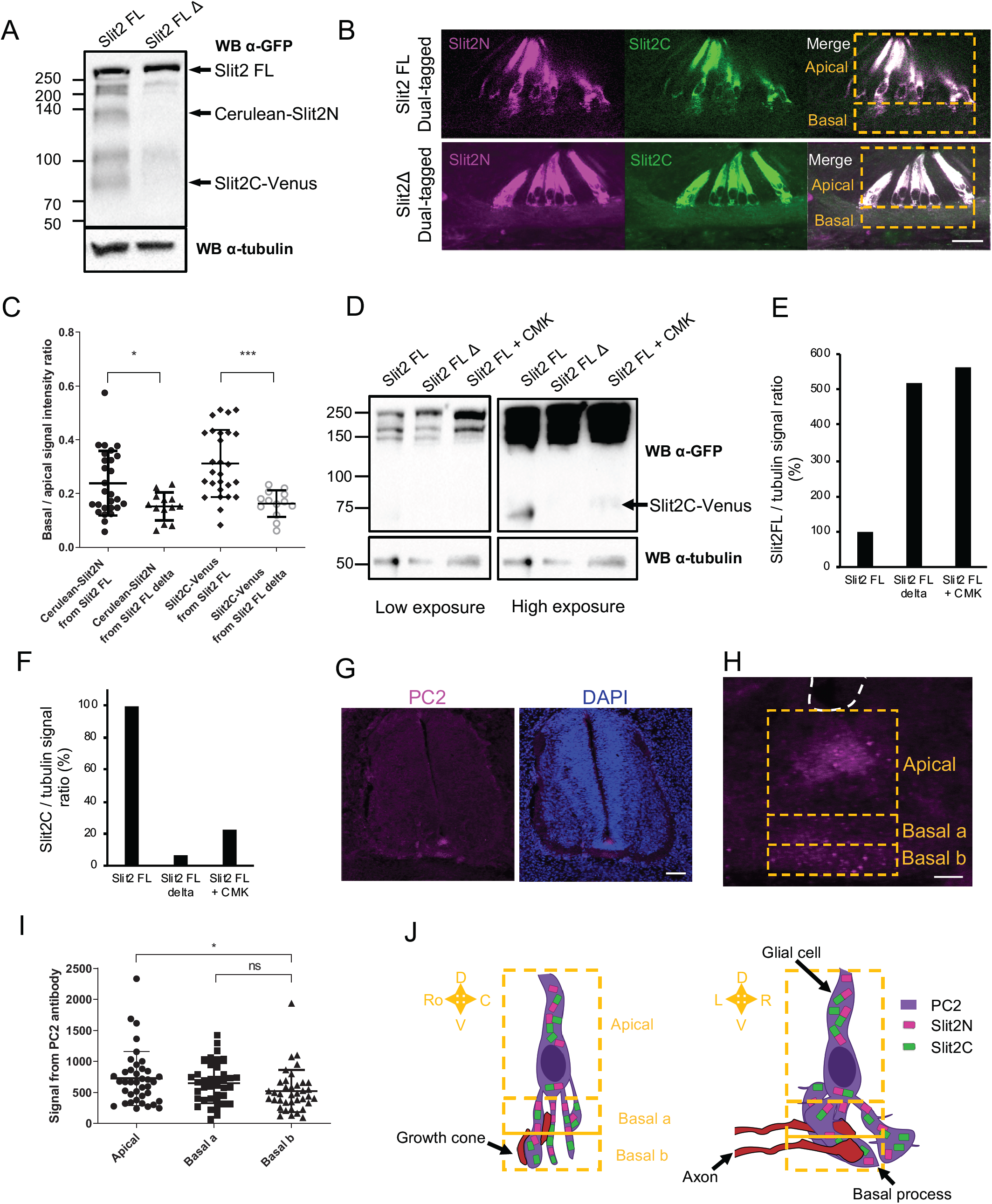
Slit2 FL cleavage plays a role in the proper diffusion of the protein and is dependent on PC2. (A) Western blot detection of Cerulean and Venus in N2a cells transfected with either dual-tagged Slit2 FL or the uncleavable dual-tagged Slit2 FL Δ. An anti-GFP antibody was used, which recognizes Cerulean and Venus, two GFP derived fluorescent proteins. The protein sizes are indicated on the left (unit: kDa). (B) Thick transverse sections of E4 chick spinal cord FP electroporated with dual-tagged Slit2 FL or dual-tagged Slit2 FL Δ (uncleavable form deprived from the cleavage site generating Slit2N and Slit2C fragments). (C) Intensity ratio of the basal compartment over the apical compartment for Cerulean and Venus in FP electroporated with dual-tagged Slit2 FL or dual-tagged Slit2 FL Δ. (D) Western blot detection of Cerulean and Venus in N2a cells transfected with either dual-tagged Slit2 FL or the uncleavable dual-tagged Slit2 FL Δ and treated with PC2 inhibitor CMK. Tubulin is used as a loading control. (E-F) Ratio of the intensity of the Slit2 FL band (E) or the Slit2C band (F) over the intensity of tubulin band. (G) Immunofluorescent labelling of E4 chick spinal cord transverse sections using an antibody targeting PC2 (left panel) and DAPI (right panel). (H) Close-up of the PC2 labelling in the FP. (I) Quantification of the PC2 labelling in the three compartments delineated by yellow dashed lines. (J) Schematic representation of FP glial cells with Slit2 FL, Slit2 fragments, and PC2 distribution as seen in a transverse section (left side) or sagittal section (right side). Data are shown as the mean ± s.d. in (C-H) and Student test has been applied. ns: non-significant, *: p < 0.05, ***: p < 0.001. Scale bars = 10µm in (B, I), 80µm in (B, G, H).

### Proteolytic processing by PC2 convertase is required for proper patterns of Slit2N and Slit2C

To decipher whether Slit2 processing contributes to the regionalization of Slit fragments in the native context, we constructed a plasmid encoding a dual-tagged Slit2 FL, fused to Cerulean in its N-terminal part and to Venus in C-terminal part, having a deletion of the Slit cleavage site (Amino-acids 110 to 118 -SPPMVLPRT-Nguyen Ba-Charvet et al, 2001). We verified by western-blot that the expression of this Slit2 FL Δ in neuronal N2a cells was not compromised. As expected, using an anti-GFP antibody (recognizing both Cerulean and Venus), the Cerulean-Slit2N and Slit2C-Venus fragments were detected in the Slit2 FL condition but not in the Slit2 FL Δ condition (Figure 6A). Dual-tagged Slit2 FL and Slit2 FL Δ were then electroporated in chicken embryos and transversal sections were analyzed by confocal microscopy. Notably, preventing Slit2 cleavage resulted in strong depletion of Slit2 protein in the basal compartment in which commissural axons navigate (Figure 6B, C). Thus, Slit2 FL protein might not be addressed to the basal processes, rather distributed in the apical compartment of FP glia cells, where it might locally be processed, generating fragments that are subsequently deposited on the basal processes. Thus, this confirmed that the cleavage is determinant for proper localization of Slit2 fragments to the axon navigation path. While the protease responsible for Slit2 cleavage is still unknown in vertebrates, the pheromone convertase Amontillado has been shown to cleave Slit during drosophila muscles and tendons development (Ordan and Volk 2016). Amontillado homolog in vertebrates is PC2, a proprotein convertase (PC) involved in the activation of endocrine peptides (Smeekens and Steiner, 1991). We treated N2a cells transfected with dual-tagged Slit2 FL with Chloromethylketone (CMK), a PC2 inhibitor, and found that it resulted in a loss of the Slit2C-Venus fragment and an accumulation of the FL form, as also observed in N2a cells transfected with Slit2 FL Δ. (Figure 6D-F). Thus, PC2 might either directly or indirectly be involved in Slit2 cleavage.

Transverse sections of E12.5 embryos were immunolabeled with an anti-PC2 antibody to examine PC2 expression at the spinal cord midline. We observed that PC2 was expressed in the FP, exhibiting a discrete and almost exclusive expression in this spinal cord territory (Figure 6G). We quantified the distribution of PC2 in the apical, basal a and b compartments and found that it is enriched in the apical compartment with modest expression in the basal compartments (Figure 6H, I). PC2 is thus properly expressed at an appropriate timing and location to mediate the cleavage of Slits and participate in the setting of their distribution patterns in the FP navigation path (Figure 6J)

### Y1815F mutation releases SlitC-mediated constrains imposed by the basal processes, resulting in abnormally plastic and exploratory growth cones during FP navigation

Our analysis revealed that SlitC does not accumulate in specific regions, nor it appeared to form a sharp gradient within the FP. Rather it distributes in clusters that decorate a tight mesh of ramified basal processes which spans the entire commissural path. Guidance cues have versatile effects, that largely depend on the context in which cues are presented (Nawabi and Castellani, 2011). The repulsive activity attributed to Slits in the context of commissural axon navigation was deduced from both *ex vivo* culture assays in which cues were delivered as soluble molecules, and interpretation of phenotypes resulting from *in vivo* manipulations (Zou et al, 2000; Long et al, 2005; Delloye-Bourgeois et al, 2015). SlitC spots could mediate a repulsive effect. Alternatively, they could act differently, for example by providing a positive signal at each contact, stabilizing the growth cone forward trajectory. Both modes of action should be reflected in the aspect of the growth cones. Thus, to get further insights, we thought to compare the morphologies of growth cones expressing PlxnA1^WT^ or PlxnA1^Y1815F^ during their navigation of the FP. E2 Chicken embryos were electroporated with plasmids encoding pHluo-PlxnA1^WT^ and pHluo-PlxnA1^Y1815F^. Open-books were mounted, immunolabeled with anti-BEN antibody to delineate FP cells and observed using confocal microscope. Within the BEN^+^ expression domain, lying in the dorso-ventral dimension below the soma territory, in which commissural axons navigate, two classes of growth cone shapes were observed: those having an oblong shape, modestly enlarged compared with the adjacent axon segment, and those having a more complex morphology, with visible protrusions and being enlarged of more than 2 times that of the axonal tract size (Figure 7A). We observed that while in the pHluo-PlxnA1^WT^ condition the proportion of complex growth cones was very low, it was increased by 3-fold in the pHluo-PlxnA1^Y1815F^ condition, reaching more than 15% (Figure 7B). Strikingly, these differences of growth cone morphologies were also very apparent in open-books labeled with Atto-647N-conjugated GFP nanobodies observed using STEP microscopy (Figure 7C).

**Figure 7:**
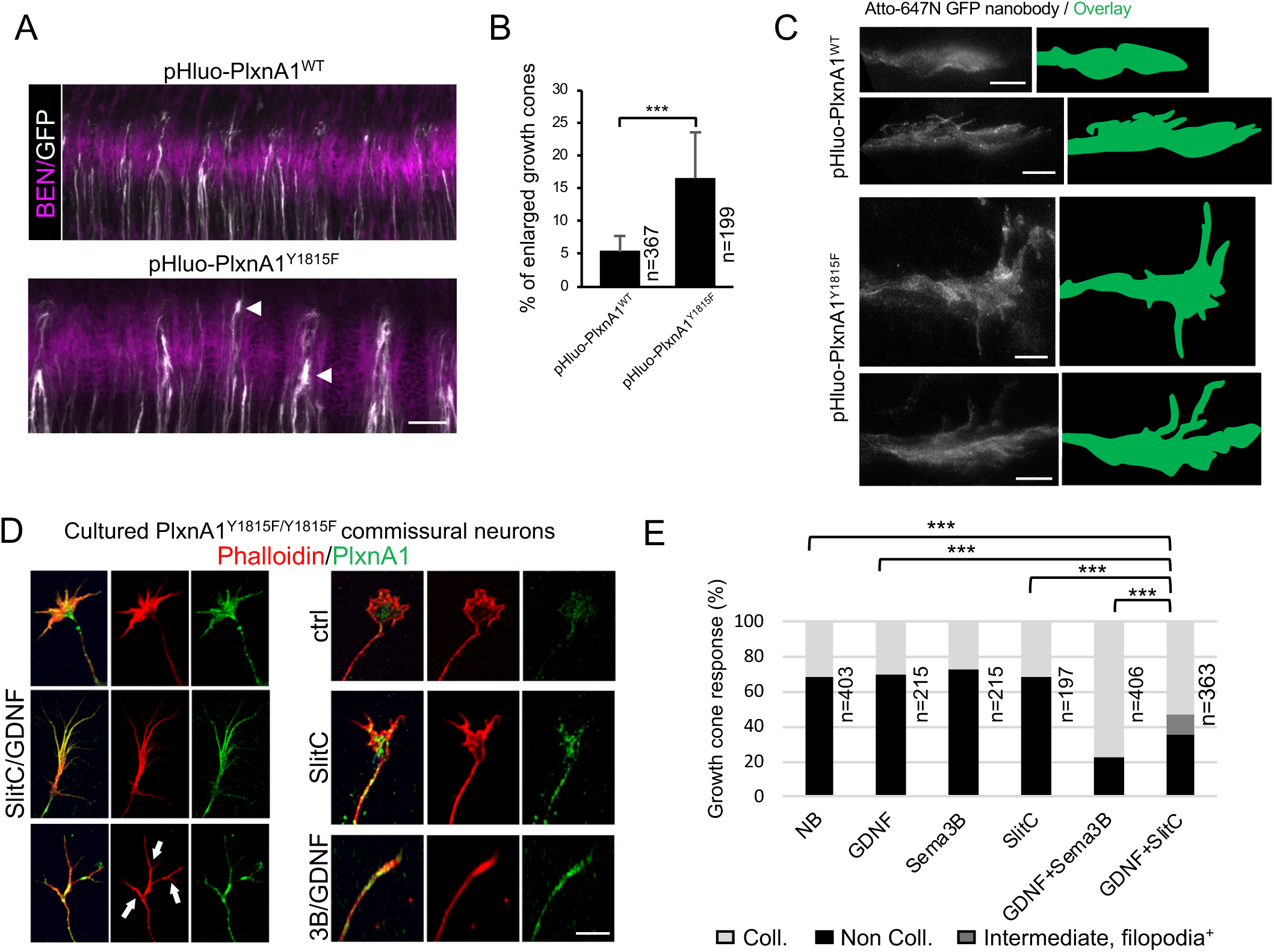
PlxnA1Y1815F mutation confers to commissural growth cones navigating the FP a complex morphology. (A) Microphotograph of spinal cord open-books from chick embryo electroporated with pHluo-PlxnA1^WT^ or pHluo-PlxnA1^Y1815F^, and immunolabeled with anti-BEN antibody and Atto-647 GFP nanobodies. BEN labeling delineates the FP. Note the complexification of growth cone morphology in the PlxnA1^Y1815F^ condition, with enlarged size and the presence of filopodia, compared with the PlxnA1^WT^ one. (B) Comparative analysis of the proportion of complex growth cones observed after PlxnA1^WT^ and PlxnA1^Y1815F^ electroporation (PlxnA1^WT^, N = 5 embryos, 367 growth cones; PlxnA1^Y1815F^, N = 4 embryos, 199 growth cones). Spinal-cord open-books were subdivided in 3 fragments and observed by confocal microscopy as a dorso-ventral stack (27 stacks). A representative image for each condition is shown. Data are shown as the mean ± s.e.m, Student test has been applied, ***: P-value p < 0.001. (C) Microphotographs of representative growth cones from STEP microscopy of atto-647N-GFP nanobodies-labeled chick open-books. Note the complexity of PlxnA1^Y1815F^ growth cones compared to PlxnA1WT ones. Scale bar: 5 µm. (D) Microphotographs of cultured PlxnA1^Y1815F/Y1815F^ commissural neuron illustrating representative growth cone morphologies exposed to different treatments. SlitC-GDNF-treated growth cones that were not collapsed exhibited a variety of shapes, some having a spread aspect, other long filopodia or forming a terminal branches pattern as shown with white arrows. (E) Histogram depicting the proportion of collapsed growth cones in the different experimental conditions (NB, N = 403; GDNF, N = 215; Sema3B, N = 215; SlitC, N = 197; GDNF+Sema3B, N = 406; GDNF+SlitC, N=363). The intermediate condition represents growth cones with a shrinked central domain but still having filopodia as shown in (D) with white arrows. Note that the presence of the Y1815F mutation in PlxnA1 reduced the growth cone collapse response, compared with Sema3B. Chi^2^ test has been applied, ***: P-value p < 0.001. Scale bars: 5µm in (C,) 15µm in (D), 50µm in (A).

These findings were evocative of SlitC acting as a repulsive molecule for commissural growth cones navigating the FP. Such repulsive effect was also suggested by previous *in vitro* experiments, reporting that soluble SlitC and Sema3B both had a collapsing effect in condition when cultured commissural neurons were exposed to the FP signal GDNF, which induces cell surface sorting of PlxnA1 receptor (Charoy et al, 2012; Delloye-Bourgeois et al, 2015). In contrast, commissural neurons over-expressing PlxnA1^Y1815F^ were significantly less sensitive to this SlitC effect, while their collapse response to Sema3B was preserved (Delloye-Bourgeois et al, 2015). We examined the behaviors of commissural neurons isolated from PlxnA1^Y1815F/Y1815F^ embryos when exposed to SlitC and Sema3B, in the absence or presence of GDNF. As expected from our previous over-expression experiments, the capacity of SlitC to collapse PlxnA1^Y1815F/Y1815F^ commissural growth cones was significantly reduced, compared to that of Sema3B. Interestingly, beyond the binary collapsed/non collapsed classification, we noted in the SlitC/GDNF condition, growth cones having atypical shapes. Some had long filopodia, or appeared contracted with numerous reminiscent filopodia or were also visibly spread (Figure 7D, E). This suggested that under normal condition, SlitC/plxnA1 signaling might negatively regulate the complexity of growth cone morphologies.

How could be translated this increased complexity of PlxnA1^Y1815F+^ commissural growth cones into aberrant axon trajectories? To address this question, we recorded the FP navigation in open-books at high frequency of time-lapse image acquisition. pHluo-PlxnA1^WT^ and pHluo-PlxnA1^Y1815F^ plasmids were electroporated in chick embryos and their spinal cords mounted in open-book. Images were taken with a time interval of 8 minutes, that enabled monitoring the growth cones over a period of time covering the FP navigation with manageable toxicity (see method). Individual growth cones were traced over time and scored according to two types of behavior: “straight” (growth cones that maintained forward trajectory), and exploratory (growth cones that deviated from their straight axis) (Figure 8A). Strikingly, while exploratory behavior was observed in only 15% of the pHluo-PlxnA1^WT+^ population, this level reached 59% in the PlxnA1^Y1815F+^ one (Figure 8B, movies S13-S16). Exploratory growth cones could either present a split aspect or could be “turned” (deviated from their straight axis, orienting in rostral or caudal directions). While turned growth cones were found equally represented in both conditions, split PlxnA1^Y1815F+^ growth cones growth cones were much more frequent than PlxnA1^WT^ ones, representing 49% of the observed cases (Figure 8C, D). Moreover, exploratory behaviors were not appearing after midline crossing or after FP exit, but were present from the onset of the FP navigation. To refine our analysis, we measured the width of individual growth cones over time, as a read-out of morphological plasticity and exploratory behavior. We found that PlxnA1^Y1815F+^ growth cones were significantly larger than those expressing PlxnA1^WT^. In addition, their increased exploratory behavior was also reflected in the higher variations of width over time when compared with PlxnA1^WT+^ growth cones (Figure 8E-F).

**Figure 8:**
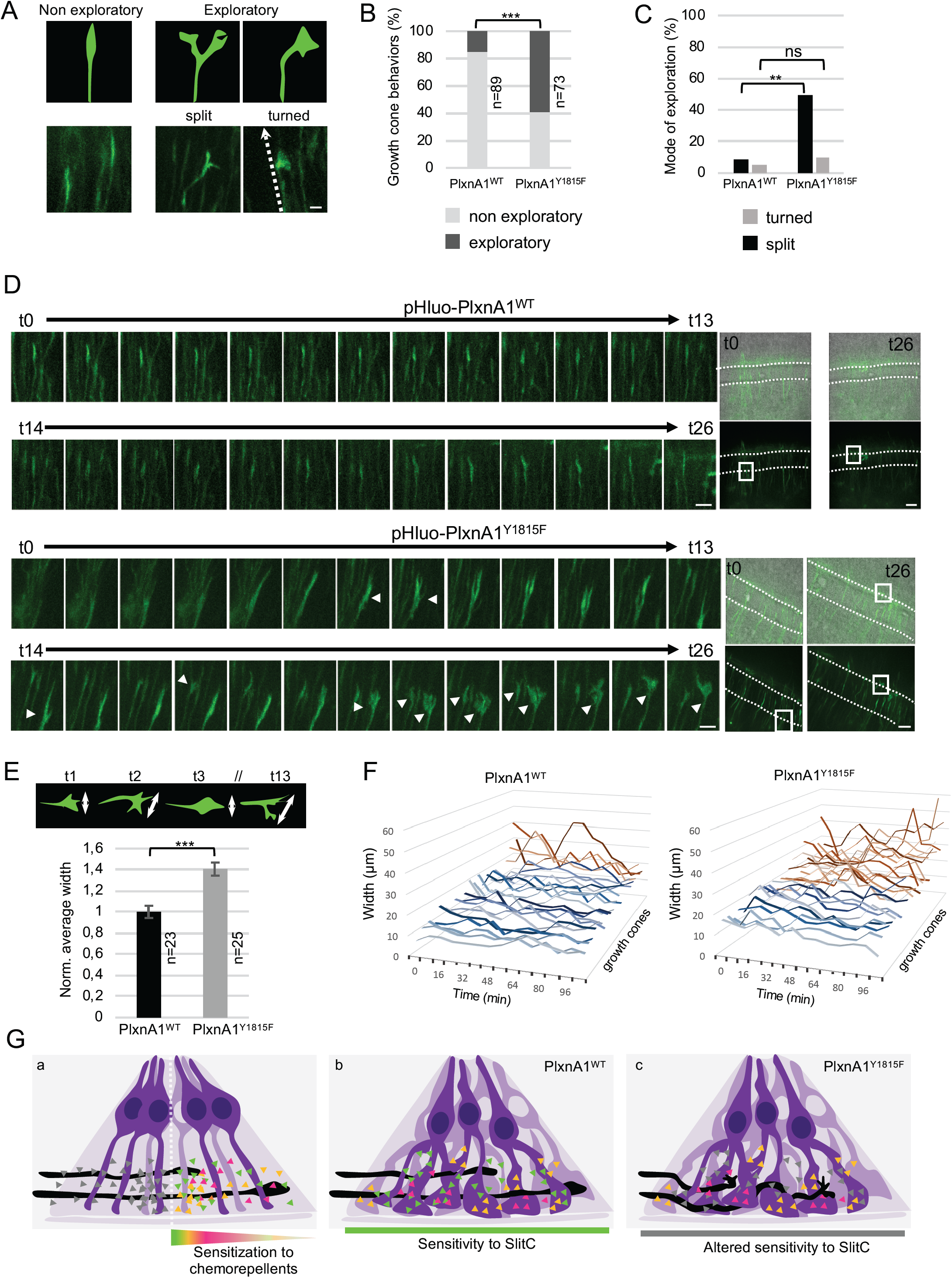
PlxnA1^Y1815F^ commissural growth cones have an increased exploratory behavior exerted through morphological split. (A) Analysis of growth cone behaviors with fast time-lapse sequences. Schematic representation of growth cone categories and microphotographs of representative growth cones. (B) Histogram of the quantification of exploratory growth cones in PlxnA1^WT^ and PlxnA1^Y1815F^ open-books (pHluo-PlxnA1^WT^, N = 5 electroporated embryos, 89 growth cones; pHluo-PlxnA1^Y1815F^, N = 3 electroporated embryos, 73 growth cones). Chi^2^ test has been applied, ***: P-value p < 0.001. (C) Histogram depicting the mode of exploration adopted by PlxnA1^WT^ and PlxnA1^Y1815F^ growth cones. The percentage were calculated over the total growth cone population (pHluo-PlxnA1^WT^, N = 5 electroporated embryos, 89 growth cones; pHluo-PlxnA1^Y1815F^, N = 3 electroporated embryos, 73 growth cones). Chi^2^ has been applied between non exploratory and split populations (**: P-value <0.01), and between non exploratory and turned (ns: non-significant). (D) Time-lapse sequences of individual growth cones navigating the FP. The right panels show growth cone positions at time 0 and time 26. Note the simple, straight, and invariable aspect of the growth cone in the PlxnA1^WT^ condition. In contrast, the morphology of the PlxnA1^Y1815F^ growth cone is much complex and varies over time. Time interval: 8 min. (E) Quantification of growth cone width navigating the FP in PlxnA1^WT^ and PlxnA1^Y1815F^ open-books, showing significant enlargement of PlxnA1^Y1815F^ growth cones compared with PlxnA1^WT^ ones (pHluo-PlxnA1^WT^, N = 4 electroporated embryos, 23 growth cones; pHluo-PlxnA1^Y1815F^, N = 3 electroporated embryos, 25 growth cones). Data are shown as the mean ± s.e.m, Student test has been applied, ***: P-value p < 0.001. (F) Histograms of individual growth cone width, in µm, from t = 0 min to t = 122 min showing increased variations of width over time in the PlxnA1^Y1815F^ condition compared with the PlxnA1^WT^ one (pHluo-PlxnA1^WT^, N = 4 electroporated embryos, 23 growth cones; pHluo-PlxnA1^Y1815F^, N = 3 electroporated embryos, 25 growth cones). (G) Current and proposed novel model of the mechanisms ensuring proper midline crossing of spinal cord commissural axons. (a) In the current model, commissural axons acquire sensitivity to FP repellents after midline crossing, and whose graded expression drive their growth direction towards the FP exit. (b) In the proposed model, commissural axons perceive SlitC from their entry in the FP. The ligands are deposited on a mesh of basal process ramifications elaborated by the FP glia. Repeated contacts with the ligands prevent the growth cones from exploring the 3 dimensions of the territory, and are maintained straight. This mechanism is ensured by functional PlxnA1-SlitC signaling. (c) Growth cones expressing PlxnA1^Y1815F^ receptor failing to properly respond to SlitC elaborate a complex peripheral structure that enables them actively exploring their local environment, which disorganizes their trajectory, leading to drastic cases to back turning. Scale bars: 15µm in (A, D left), 50µm in (D right).

Overall, these results support a model whereby SlitC patches presented by the basal processes network spanning the entire FP navigation path generate a straight trajectory of the growth cones by continuously limiting their exploration potential through inhibitory/repulsive repetitive contacts.

## Discussion

Our results establish that commissural axons navigate the FP in a complex environment, composed of ramified FP basal processes with stereotypic spatial organization. These basal structures are decorated with segregated Slit2C, Slit2N, and Sema3B clusters, forming a mesh that forces multiple and reiterated contacts with the navigating growth cones. Through analysis of a knock-in mouse model baring a PlxnA1 mutation specifically altering Slit2C responses and a range of *ex vivo* and imaging analysis, we show that the Slit2C-PlxnA1 signaling is indispensable for the maintenance of forward-directed commissural axon trajectories. We also identify Y1815 of PlxnA1, highly-conserved across PlxnA family members, as a key residue for proper distribution at the cell membrane. Finally, our data do not support that midline recrossing is prevented through gain of sensitivity to repellents after midline crossing. On the contrary, they show that repulsive forces generate a forward polarity of commissural axon growth starting from the FP entry and being active over the entire FP navigation. We propose an alternative view of the mechanism ensuring FP crossing, which is to limit continuously over the FP navigation the exploration activity of commissural growth cones (Figure 8G).

### Specific sub-cellular architecture of the FP glia conforms the presentation of FP ligands to the navigating commissural axons

The mode of presentation of the FP ligands to the navigating axons has remained largely ignored, although it is generally admitted that it is determinant for the final outcome of guidance cues. Hence, despite extensive studies of spinal cord and midbrain midline crossing, very little is known on the tridimensional organization of the FP. Early electronic microscopy images revealed close contacts between FP glia surfaces and axons, and suggested complex shapes of basal processes (Yaginuma et al. 1991; Campbell and Peterson, 1993; Okabe et al. 2004). To which extent, and if so how, the spatial organization of the FP glia generates specific ligand patterns has been left open. Our study sheds novel light on this issue. Analysis of ligands distribution with fluorescent reporters whose expression was specifically driven in FP cells and immunolabeling of the FP-specific marker, BEN, allowed us to reconstruct in 3D the FP glia morphology. A particularly striking observation was the asymmetric apico-basal polarity of the FP cell, with a single apical anchor and an enlarged basal process subdivided into several oblong pillars. In addition to this morphological complexity, we found that the basal ramified structures have a stereotypic spatial position. Their larger sides align along the left-right and rostro-caudal axes, parallel narrow corridors in which commissural axons navigate. Finally, in the dorso-ventral axis, ramifications of the main pillars also provide to the axons a net-shaped roof. Notably, this sub-cellular architecture of the FP glia provides a physical frame for the presentation of the ligands. We found that SlitC, SlitN, and Sema3B have resembling distribution patterns, prominently deposited in columnar clusters covering the ramified basal processes of FP glia cells. Nevertheless, our analysis using dual tagged Slit2 construct indicated that SlitC and SlitN clusters occupy physically distinct locations.

The molecular scaffold that localizes the ligands in such a way is still to be explored. Extracellular matrix components and their interactors are well acknowledged for their critical role in setting molecular patterns (Walter et al, 2018). Interestingly in the *zebrafish* developing brain, apical end feet of radial glial cells extend at the surface of the tectum, organizing through collagen IV the lamination of the neuropil by exposing Slits and heparan sulfate proteoglycans to axon terminals (Xiao et al, 2011). Morphogenesis of the *drosophila* heart tube lumen was found to rely on polarized localization of the collagen XV/XVIII orthologue Multi-Plexin forming a macro-complex with Slit (Harpaz et al, 2013). In the context of midline crossing, loss of the heparan sulfate carrier (Heparan sulfate proteoglycan) Syndecan results in modification of Slit distribution and activity in *drosophila* (Johnson et al, 2004). In the vertebrate spinal cord, dystroglycan was identified as a Slit binding partner, localizing the proteins in the basal lamina through non-cell autonomous action (Wright et al, 2012; Lindenmaier et al, 2019). Consistently, we detected the presence of SlitC, whose sequence contains dystroglycan binding motif, in the basement membrane, as well as that of SlitN, possibly sequestrated by additional components of the FP basal lamina. For example, proteolysis of the extracellular matrix protein F-spondin, reported to contribute to commissural axon guidance (Burstyn-Cohen et al, 1999), was found to generate two fragments, among which one is deposited at the basement membrane (Zisman et al, 2007). Patterning of guidance molecules by morphological specificities of producing cells was also nicely exemplified on Netrin-1. Synthesized by progenitor soma in the ventricular zone of the neuroepithelium, Netrin-1 was proposed to be transported on or within the basal process for relocalization at the basal membrane, where it might serve as a growth-promoting substrate for pre-crossing commissural axons (Varadarajan et al, 2017).

Beyond the molecular components localizing the ligands, our analysis of Slit fragments distribution also revealed that Slit cleavage is a pre-requisite for the detection of Slits in the commissural axon navigation path, since uncleavable Slit2 almost exclusively localized in the FP apical domain. This suggests that, unlike in *drosophila* midline guidance for which Slit processing was reported to be dispensable (Coleman et al, 2010), under their full-length form, Slits likely play limited role, if not no role at all, in the guidance of spinal cord commissural growth cones. We also found that SlitC, when synthesized as a cleaved fragment, was significantly more prone to basement membrane deposition than its equivalent form resulting from endogenous processing. These findings support the intervention of a series of post-translational processes to properly present the ligands to the commissural axons, in particular those controlling the addressing of full-length Slit in the FP glia sub-cellular compartments, as well as those regulating the protease action. This is consistent with previous studies of the process of muscle anchorage to tendons in *drosophila*, that reported the importance of Slit processing for subsequent distribution of the fragments and proper functional outcome (Ordan and Volk 2015).

### Commissural axon navigation is driven by restriction of growth cone exploration resulting from reiterated contacts with ligands deposited on the FP basal processes

A puzzling question of commissural axon navigation across the FP relates to the mode of action by which FP cues exert their effect. Proper FP crossing is thought to rely on a balance of positive and negative forces that support growth cone attachment, motility and direction (de Ramon Francàs et al, 2017). Early experiments conducted in the chicken embryo demonstrated that contacts engaging various Ig Superfamily Cell Adhesion Molecules expressed by growth cones and glia cells control the entry in the FP (Stoeckli et al, 1997; Fitzli et al, 2000). Complementarily, growth-promoting effect exerted by the Stem Cell Factor (SCF) under its transmembrane isoform acting via its KIT receptor was shown to be switched on after midline crossing to facilitate the growth towards the FP exit (Gore et al, 2008). Finally, studies of midline crossing in *drosophila* inspired the view that in vertebrates as well commissural axons must acquire sensitivity to local repellents after midline crossing, in order to get directional information for exiting the FP and for not turning back to the ipsilateral side (Evans and Bashaw, 2010). Consistently, differential sensitivity of commissural growth cones to FP cues before and after FP crossing was reported in numerous experimental paradigms (Zou et al, 2000; Nawabi et al, 2010; Delloye-Bourgeois et al, 2015). Nevertheless, in vertebrates, the physical territory of midline navigation, the FP, is much larger and complex than that in invertebrates, which might impose particular constraints and require specific guidance forces that remain uncharacterized. Indeed, the exact mechanisms by which SlitC, SlitN, and Sema3B exert their action in the FP are unknown, although they are all considered as chemorepellents. However, this has yet to be demonstrated since they were studied using only *ex vivo* paradigms that neither recapitulated the architecture of the FP navigation path, nor the coincident information that the growth cones receive from their local environment when navigating in their native context (Zou et al, 2000; Nawabi et al, 2010; Delloye-Bourgeois et al, 2015). Functional outcomes of guidance signals are indeed highly versatile and shown to depend on many intrinsic and extrinsic variables (Nawabi et al, 2011). *In vitro*, context-dependent switch from attraction to repulsion and vice-versa were evidenced for a large range of guidance molecules including Netrins, Semaphorins, and Slits (Höpker et al, 1999; Castellani et al, 2000; Nguyen-Ba-Charvet et al, 2001; Castellani et al, 2002; Ma et al, 2007). *In vivo*, although considered as repellents, Sema2a and Sema3B were recently reported to promote midline crossing in *drosophila*, acting as chemoattractant via Sema1a (Hernandez-Fleming et al, 2017).

Our findings provide the first evidence that specific FP guidance forces ensure commissural axons a continuous straight trajectory, from the FP entry to the exit. With our analysis of PlxnA1^Y1815F^ model and live imaging paradigms, we show that SlitC brings a major contribution to this guidance mechanism, acting by restricting the exploratory capacity of commissural growth cones. This function is served by the particular morphology of the FP glia cells that might impose repeated contacts of the growth cones with spots of ligands spanning the navigation path. Our functional analyses are all consistent with SlitC mode of action restricting the formation or the stabilization of peripheral growth cone protrusions to constrain them in a non-exploratory state, thus favoring forwards growth direction. Such effect is also reflected in the shape of wild-type crossing growth cones, which we observed are rostro-caudally thin and dorso-ventrally elongated. Notably, PlxnA1^Y1815F+^ growth cones have a much more complex morphology than WT growth cones, and they also sense the environmental content much more actively. In recent work, axon growth was modeled using microcontact printing culture devices developed to examine individual growth cone responses to dots of guidance molecules (Ryu et al, 2018). Axons were challenged to grow on micropatterned surfaces consisting in uniform Sema3F substrate interrupted by permissive dots. They were found able to efficiently extend in such surfaces, having, in addition, a straight trajectory resulting from jumping from one permissive dot to the next one. Interestingly, perturbing the distance between dots or their size modified the shape of the axons by disorganizing their cytoskeleton and their trajectory, inducing curved growth patterns (Ryu et al, 2018). Thus, a “salt and pepper” context as in the FP with repulsive ligands localized in spots and intercalated with permissive regions might build an appropriate topographic environment for limiting the possibilities of growth deviations.

Such constrain of growth cone exploration might also be required for counteracting other local coincident guidance forces that would disturb axon trajectories during FP crossing. In our recent work, we found in live imaging that the exploration is increased when growth cones start navigating the second FP half, although we observed they still keep a clear straight growth (Pignata et al, 2019). This exploratory behavior could reflect an increasing sensitivity to FP-derived SHH and WNT rostro-caudal gradients that were found to drive the longitudinal turning after the crossing (Zuñiga and Stoeckli, 2017). The exact timing at which the axons become sensitive to these gradients is unclear. In Robo3 knockout embryos, commissural axons totally fail to cross the FP but turn longitudinally in the ipsilateral side. This argues that non-crossing commissural axons can get sensitized to longitudinal gradients, which also reflects the need for guidance constrains to counteract premature turning during the FP navigation (Friocourt and Chedotal, 2017). One function of PlxnA1/SlitC signaling would thus be to prevent premature longitudinal turning until FP exit.

### Tyrosine 1815 mutation alters the membrane mobility of PlxnA1 receptor

Tyrosine phosphorylation is a pivotal modification by which receptors can acquire key specific dynamics, trafficking, or signaling properties. Plxn cytoplasmic domain contains several tyrosine residues, some of them being phosphorylated during the signaling cascade downstream of Semaphorin ligands (Tamagnone *et al*, 1999). Apart from PlxnB (Swiercz et al, 2009), the characterization of the molecular events triggered by specific tyrosine phosphorylation and identification of kinases targeting the different residues is only beginning. Recently, two highly-conserved residues were shown as major targets of Fyn, a kinase known to phosphorylate PlxnA1 downstream of Semaphorins, and their phosphorylation is required for *zebrafish* eye development (St Clair et al, 2018). All Plxns share a set of thirteen tyrosine, but interestingly the Y1815 is restricted to the PlxnA subfamily (Franco and Tamagnone, 2008). Consistently, we showed that PlxnAs interact with SlitC while PlxnBs do not (Delloye-Bourgeois et al, 2015). The present study shows that mutating the Y1815 residue strongly impacts on the distribution of PlxnA1 at the cell membrane. First, we observed in mouse commissural axons navigating the FP in PlxnA1^-/-^ open-books that the PlxnA1^Y1815F^ cell surface pool expanded to the adjacent axon shaft, whereas it was much more restricted to the growth cone compartment in the WT context. Second, when expressed in chicken open-books for STED imaging, PlxnA1^Y1815F^ was observed to distribute at the membrane of the entire growth cone, not accumulating at the front as normally observed for PlxnA1^WT^ (Pignata et al, 2019). Third, FRAP experiments revealed an increased membrane mobility of the PlxnA1^Y1815F^ receptor pool, compared with the WT one, with a pattern of fluorescence recovery suggesting that exocytosis-mediated sorting is in contrast unaffected by the mutation. Overall this raises the interesting possibility that PlxnA1 baring Y1815F mutation lacks an interactor whose function in stabilizing the receptor at the growth cone membrane is indispensable for SlitC signaling. Structural analysis established the ability of PlxnA1 to engage its extra-cellular domain in dimeric complexes (Janssen *et al.*, 2010; Kong *et al.*, 2016). Thus, Y1815 could be required for allowing the formation of macro-complexes of PlxnAs at the growth cone membrane. Neuropilins are unlikely involved since we found they are dispensable for SlitC-PlxnA1 binding and SlitC responses (Delloye-Bourgeois et al, 2015). Heparan sulfate proteoglycans are possible candidates since they bind PlxnA1 (Delloye-bourgeois et al, 2015). Y1815 could also be involved directly or indirectly for the receptor to recruit scaffolding proteins and intracellular partners. For example, a recent study reported a novel interface between PlxnB2 and the PDZ domain of PDZ-RHO-GEF, whose activation downstream of Semaphorin binding increases GTP-bound RHOA activity, that maps at the C-terminus side engaging an amino-acid sequence close to the 1815 residue (Pascoe et al, 2015). Further studies are needed to elucidate the contributions of Y1815 residue to the PlxnA1 biological activity.

### Is PlxnA1-SlitC signaling the only player of the guidance program generating straight trajectory across the FP?

*In vivo* re-crossing phenotypes have been observed when PlxnA1, Slit1-3, but not Robo1/2 or Sema3B, are inactivated and we show here that it is manifested when PlxnA1 is prevented to properly mediate SlitC activity. This does not exclude that the Slit/Robo signaling also contributes to this function. The lack of recrossing in context of Robo1/2 loss could be due to the specific temporal and spatial pattern of Robo1 receptor at the growth cone surface. According to our previous work (Pignata et al, 2019), growth cones navigating the first FP half express PlxnA1 but not yet Robo1, while in the second FP half, they express both. Robo2 was found to be sorted only during the post-crossing longitudinal navigation (Pignata et al, 2019). Thus, PlxnA1 and Robo loss cannot have the same observable outcome, whether or not both receptors have similar functions. PlxnA1 deletion results in lack of Slits and Sema3B receptors when the growth cones enter the FP, whereas Robo deletion manifests itself later when growth cones already completed half of their navigation through the FP, and have a preserved PlxnA1-SlitC signaling to navigate the second FP half. Moreover, recrossing is likely to be the more drastic growth cone behavior resulting from abrogation of forces counteracting rostro-caudal gradients. Interestingly in Robo1/2^-/-^ embryos, cases of commissural axons aberrantly oriented within the FP were reported (Long et al, 2004). As well in PlxnA1^-/-^ embryos, premature turning in the FP was also frequently observed (Nawabi et al, 2010; Delloye-Bourgeois et al, 2015). Such phenotypes could well be interpreted as resulting from alleviation of straight growth constrains. The equally plausible alternative scenario would be that Slit2C-PlxnA1 signaling does indeed specifically ensure the straight growth of commissural axons in the FP, through specific downstream signaling. In support, we found that PlxnA1 and Robo1 are not only sorted at different timing but also in different growth cone compartments, with PlxnA1 enriched at the front and Robo1 at the rear, thus able to generate spatially distinct functional outcomes (Pignata et al, 2019).

Whatever the case, our study raises a novel guidance model of midline crossing that might represent a general mechanism for the navigation at choice points. Limiting growth cone exploration through short-range signaling would efficiently allow inhibiting premature directional changes elicited by coincident guidance cues, until intermediate target navigation is completed.

## Materials and Methods

### Generation of the PlxnA1^Y1815F^ mouse line and genotyping

The PlxnA1^Y1815F^ mouse line was generated by the Mouse Clinical Institute (Strasbourg, France). Mice were back-crossed to obtain PlxnA1^Y1815F^ in a C57/BL6 background. Mice were hosted either in SPF (ALECS SPF, Lyon, France) or conventional (SCAR, Lyon, France) animal facility with *ad libitum* feeding. This study was covered by a Genetically Modified Organisms approval (number 561, French ministery for Research) and a local ethical comity (CECCAPP, Lyon). Genotyping was performed on dissected tails lysed in NaOH 1M solution for 25 min at 95°C and digested overnight with Proteinase K at 56°C with the following primers 5’-CTT ATA GAT CTA GAC AGG CAG GGA GAC CAT-3’ and 5’-CGG TTG TCT TCT CGA GTA TCA CAC TCC TA-3’. The PCR kit used was FastStrat PCR Master (Roche, 04710436001). The amplification product from mutated allele has a 341bp size and the one obtained from a wild-type allele is 262bp. Genotyping of PlxnA1-/- was done as in Yoshida et al. (2006).

### Molecular biology

FL mouse pHluo-PlxnA1 was generated by introducing in Nter the coding sequence of the pHluo cloned from a vector encoding GABA A pHluo-GFP (Jacob et al., 2005). FL mouse pHluo-PlxnA1^Y1815F^ was obtained by directed mutagenesis using the In-fusion HD Cloning Plus Kit (638909, Ozyme) with the following primers: 5’ AGG TAC TTT GCT GAC ATT GCC and 5’ GTC AGC AAA GTA CCT TTC AAC. The pH dependency of the fluorescence was validated as in Pignata et al (2019).

Dual-tagged Slit2 was designed using the human Slit2 sequence (NM_001289135.2) and ordered from Genscript Biotech. Briefly, Cerulean was inserted after Slit2 signal peptide (MRGVGWQMLSLGLVLAILNKVAPQACPA) and Slit2 FL sequence. Venus was fused to the C-terminal part of Slit2. Dual-tagged Slit2 Δ construct was obtained by Quickchange mutagenesis procedure (Agilent), deleting 9 amino acids (SPPMVLPRT) at the cleavage site. The primer used for the mutagenesis were 5’-CTT GTT CTG TGA GTTT AGC CCC TGT GAT AAT TTT G-3’ and 5’-CAA AAT TAT CAC AGG GGC T AAA CTC ACA GAA CAAG-3’. Both constructs were cloned into a PCAGEN vector using NotI and EcoRV digestion. The Hoxa1-GFP plasmid was a kind gift of Esther Stoeckli.

The fusion of Slit2N and Slit2C with GFP was performed in order to express the fusion proteins resulting from the cleavage of Slit2 Dual-tagged construct: GFP-SlitN (GFP in Nter) and SlitC-GFP (GFP in Cter). The fusion of Slit2N with GFP was done by cloning the Slit2 signal peptide, the eGFP and the Slit2N sequences in a pCAGEN backbone using the In-Fusion HD cloning Plus Kit (638909, Ozyme) with the following primers : 5’-GCA AAG AAT TCC TCG AGG ATA TCA TGC GCG GCG TTG GCT GG-3’ and 5’-TGC TCA CGT TAA CCG CCG GGC ACG CCT G-3’ for the signal peptide; 5’-CCC GGC GGT TAA CGT GAG CAA GGG CGA GG-3’ and 5’-AGC ACT GAC GCG TCT TGT ACA GCT CGT CCA TGC-3’ for eGFP; 5’-GTA CAA GAC GCG TCA GTG CTC TTG CTC GG-3’ and 5’-CTG AGG AGT GCG GCC GCG ATT TAA CGA GGG AGG ACC ATG GG-3’ for Slit2N. The CAG promoter was then replaced by the Hoxa1 promoter using In-Fusion with the following primers: 5’-GTG CCA CCT GGT CGA CGC TTC TTC TAG CGA TTA AAT C-3’ and 5’-AAC GCC GCG CAT GAT ATC CCC ACT AGT AAG CTT GGA GGT G-3’. The fusion of Slit2C with GFP was done by cloning the sequences into a pEGFP-N3 vector after the addition of KpnI and BamHI restriction sites using the following primers: 5’-cg GGTACC acc agc ccc tgt gat aat ttt g-3’ and 5’-cg GGATCC gga cac aca cct cgt aca gc-3’. The Igκ Leader peptide signal was added for correct secretion by cloning the fusion fragment into a pSecTagB plasmid using KpnI and NotI digestion. The resulting sequence was cloned into a pCAGEN vector using XhoI and NotI digestion, and the In-Fusion HD Cloning Plus Kit (638909, Ozyme) with the following primers: 5’-caa aga att cct cga gat gga gac aga cac act cct gc-3’ and 5’-ctg agg agt gcg gcc gct tac ttg tac agc tcg tcc atg cc-3’. The CAG promoter was replaced by the Hoxa1 promoter using SalI and XhoI digestion, and the In-Fusion HD Cloning Plus Kit with the following primers: 5’-gtg cca cct ggt cga cgc ttc ttc tag cga tta aat caa ag-3’ and 5’-ctg tct cca tct cga gcc cac tag taa gct tgg agg tg-3’. Math-1-mbTomato construct was described previously (Pignata et al, 2019).

### DiI staining in spinal cord open-books

Spinal cords from E12.5 murine embryos were dissected and mounted as open-books prior to fixation in 4% paraformaldehyde (PFA) for 18 hrs. DiI crystals (D3911, ThermoFisher) were inserted in the most dorsal part of one hemi-spinal cord for anterograde labeling of commissural tracts. Axon trajectories were analyzed 24 hrs later with a spinning disk microscope (Olympus X80). For each DiI crystal, a range of phenotypes can be observed. Classes were made representing the percentage of DiI crystals showing the phenotype over the total number of observed DiI crystals. Classes were assessed independently, with percentage ranging from 0% to 100%.

### Immunofluorescence labeling

Cryosections from embryos collected at E12.5 were prepared and processed for staining as in Charoy et al, (2012). For some experiments, chick embryos sections and open-book spinal cords were blocked in 6% BSA (A7906, Sigma) and 0.5% Triton (T9284, Sigma) diluted in PBS for 5 hours at room temperature. Sections were incubated overnight at room temperature with anti-PlxnA1 antibody (gift from Y. Yoshida), anti-Robo3 antibody ((1/100, R&D, AF3076); anti-L1CAM antibody (1:100, A439 Abcam 123990), anti-NgCAM antibody (1:50, 8D9, DSHB), anti-BEN (1:50, BEN, DSHB) or an anti-PC2 antibody (1:100, 3533, Abcam) in 1% BSA diluted in PBS. Alexa 488, Alexa 555 (1/500, Invitrogen) and Fluoroprobe 546 (1/400) were used as secondary antibodies.

### *In ovo* electroporation, open-book mounting and imaging

The neural tube of HH14/HH15 chick embryos was electroporated as described previously (Delloye-Bourgeois *et al.*, 2015, Pignata et al, 2019). Plasmids were diluted in PBS with Fast Green (F7262, Sigma) at the following concentration:

- 1.5mg/mL for dual-tagged Slit2 and dual-tagged Slit2 Δ. Lower concentration did not allow correct imaging with confocal microscopy.
- 1.34mg/mL for Hoxa1-Slit2N-GFP and 0.94mg/mL for Hoxa1-Slit2C-GFP. Both concentrations match the dual-tagged plasmid molarity in order to electropore the same amount of plasmid.
- 0.5mg/mL for Math1-mbTomato and 2 μg/μl for pHluo-PlxnA1 and pHluorin-PlxnA1^Y1815F^ and 0,05mg/mL for mbTomato and Hoxa1-GFP. These concentrations were selected according to our previous work (Pignata et al, 2019) in which we compared the outcome of different concentrations on the FP navigation and selected the conditions that gave the better compromise between receptor expression and ability of FP crossing. The pHluo dependency was controlled by *in vitro* cell line transfection as in Pignata et al, (2019).

The plasmid solution was injected into the lumen of the neural tube using picopritzer III (Micro Control Instrument Ltd., UK). Using electrodes (CUY611P7-4, Sonidel) 3 pulses (25V, 500ms interpulse) were delivered by CUY-21 generator (Sonidell). Electroporated embryos were incubated at 38.5°C. In ovo electroporation of floor plate cells was done on HH17/18 chick embryos as described by Wilson et al. (2012). Briefly, electrodes (CUY611P7-4, Sonidel) were placed at the thoracic level dorsally (cathode, negative electrode) and ventrally (anode, positive electrode), and 3 pulses (18V, 500ms interpulse) were delivered by CUY21 electroporator (Sonidell). Embryos at HH25/HH26 were harvested in cold HBSS (14170-088, Gibco) and the spinal cords were dissected out. They were mounted in 0.5% agarose diluted in F12 medium on glass bottom dishes (P35G-1.5-14-C, MatTek). After agarose solidification, spinal cords were overlaid with 3ml of F12 medium supplemented with 10% FCS (F7524; Sigma-Aldrich), 1% Penicillin/Streptomycin (Sigma-Aldrich) and 20mM HEPES buffer (15630-049, ThermoFischer Scientific). For section imaging, embryos were then fixed for 2 hours at room temperature in 4% paraformaldehyde diluted in PBS. For open-book imaging, spinal cords were dissected and then fixed 45 minutes in 4% paraformaldehyde diluted in PBS. For vibratome sectioning, embryos were embedded in 3% low gelling agarose (A9414, Sigma) diluted in PBS. The embryos were then sectioned in 80µm slices using Leica VT1000S vibratome.

### Mouse spinal cord electroporation and open-book mounting

E12 mice embryos were collected and fixed on a SYLGARD (Dow Corning) culture plate in HBSS medium (ThermoFisher) supplemented with Glucose 1M (Sigma-Aldrich). Injection of plasmids into the lumen of the neural tube was performed using picopritzer III (Micro Control Instrument Ltd., UK). Using electrodes (CUY611P7-4, Sonidel) 3 pulses (20V, 500ms interpulse) were delivered by CUY-21 generator (Sonidell). Spinal cords were dissected from the embryos and cultured on Nucleopore Track-Etch membrane (Whatman) for 48 hours in Slice Culture Medium (*Polleux and Ghosh., 2002*).

### Imaging and data analysis

Live imaging was performed with an Olympus IX81 microscope equipped with a spinning disk (CSU-X1 5000 rpm, Yokogawa) and Okolab environmental chamber maintained at 37°C. Image were acquired with a 20X objective by EMCCD camera (iXon3 DU-885, Andor technology). 15-30 planes spaced of 0,5-3µm were imaged for each open-book at 30-minute interval for 10 hours approximatively. To reduce exposure time and laser intensity, acquisitions were done using binning 2×2. Images were acquired using IQ3 software using multi-position and Z stack protocols. Z stack projections of the movies were analyzed in ImageJ software. The analysis of pHluo-flashes was performed from time-lapse acquisitions. In some experiments, the time interval was reduced for faster image acquisition. Time intervals of 3 minutes, 5 minutes, 8 minutes and 12 minutes were tested. At 3 and 5 minutes, the tissues were rapidly damaged. We thus selected time interval of 8 min as the better compromise between time resolution and phototoxicity.

Confocal imaging was performed with either an Olympus FV1000 with a 40x objective and zoom or a Leica TCS SP5 with a 63x objective. Deconvolution was done using the Huygens software. 3D surface reconstructions were done using the Imaris software.

### Atto647N staining and STED microscopy

Spinal cords were incubated at 38°C for 20 minutes with F12 medium supplemented with 5% FCS (F7524; Sigma-Aldrich), 20mM HEPES buffer (15630-049, ThermoFischer Scientific) and 1/100 GFP-nanobodies Atto647N. They were then rinsed 4 times with the same medium free of GFP-nanobodies and fixed at room temperature for 2 hours with PBS supplemented with 4% paraformaldehyde (PFA) and 1% BSA (A7638 Sigma-Aldrich). Open-books were observed with a STED microscope (TCS SP8, Leica). STED illumination of ATTO 647N was performed using a 633-nm pulsed laser providing excitation, and a pulsed bi-photon laser (Mai Tai; Spectra-Physics) turned to 765 nm and going through a 100-m optical fiber to enlarge pulse width (100ps) used for depletion. A doughnut-shaped laser beam was achieved through two lambda plates. Fluorescence light between 650 and 740 nm was collected using a photomultiplier, using a HCX PL-APO CS 100/1.40 NA oil objective and a pinhole open to one time the Airy disk (60mm). Images were acquired with using Leica microsystem software and a Z stack protocol. Usually 10-20 planes spaced of 0,5µm where imaged for each growth cone. The growth cones were delineated and the intensity signal was calculated using ImageJ.

### Fluorescence Recovery After Photobleaching (FRAP)

FRAP experiments were conducted on spinal cord open-books electroporated with either pHluo-receptor and mbTomato, using a Leica DMI6000 (Leica Microsystems, Wetzlar, Germany) equipped with a confocal Scanner Unit CSU-X1 (Yokogawa Electric Corporation, Tokyo, Japan) and a scanner FRAP system, ILAS (Roper Scientific, Evry, France). Images were acquired in both green and red channels using a 63X objective and an Evolve EMCCD camera (Photometrics, Tucson, USA). Growth cones located in the FP were first monitored for 12s each 3s and then bleached using a 488nm diode laser at full power. This resulted in an 85-95% loss of the signal (mean of 89%) at t=0. Fluorescence recovery was then monitored for 730s with acquisitions every 3s for 30s, then every 10s for 100s, and finally every 30s for 600s. The images were corrected for background noise, residual fluorescence right after the bleach was set to zero, and recovery curves were normalized to the fluorescence lost after the bleach. No other corrections were applied since unbleached growth cone fluorescence showed no significant decay during the acquisition period.

### Western blot

To observe Slit cleavage, N2a cells were seeded into 6-wells plates (2.5.10^5^ cells per well). 24 hours later, cells were transfected using jetprime transfection reagent (114, Polyplus transfection). 4 hours after starting the transfection, cell medium was changed and CMK (ALX-260-022, Enzo) was added to a final concentration of 100µM if needed. Two days after transfection, CMK treatment was repeated. Two hours after, cells were harvested. Whole cells extract was isolated using RIPA buffer (NaCl 150mM – Tris HCL pH7,35 50mM – DOC 1% – N-P40 1% – H_2_O) supplemented with protease inhibitor (04 693 116 001, Roche). Isolated protein concentration was determined using Bradford assay (500-0006, Bio-Rad).

Spinal cords were isolated from E12.5 embryos and dissected tissues were lyzed in RIPA buffer supplemented with protease inhibitor. Samples were analyzed in western blot using anti-PlxnA1 antibody (Gift from Y Yoshoda), anti-GFP (1:1000, 11814460001, Sigma), anti-Tubulin (1:10000, T5168, Sigma) and anti-PC2 (1:100, 3533, Abcam). Western blot quantification was performed using Image Lab4.0 software (Bio-Rad).

### Commissural neuron cultures and collapse

Dorsal spinal cord tissue was dissected out from isolated spinal cord and dissociated. Neurons were grown on laminin-polylysine-coated coverslips in Neurobasal supplemented with B27, glutamine (Gibco), and Netrin-1 (R&D) medium for 24 to 48 hrs, as in Nawabi et al, (2010). Immunolabeling was performed with anti-PlxnA1 antibody (gift from Y. Yoshida). Nuclei were stained with bisbenzimide (Promega) and actin with TRITC-phalloidin. GDNF was applied to the cultures as in Charoy et al (2012). Collapse assays were performed as in Delloye-Bourgeois et al, 2015.

### Quantification and Statistical analyses

Protein diffusion quantification: The background noise was removed by measuring it, then subtracting it in ImageJ. The electroporated zone was then divided into two different compartments along the dorso-ventral axis. The glial cells’ apical feet and cellular bodies were included in the apical compartment. The axons path was divided in two, the most apical compartment being basal 1, the most basal compartment being basal 2. The mean intensity of Cerulean, Venus or GFP was measured using ImageJ in each compartment. The mean intensity was then normalized by the mean intensity in the apical compartment.

Pearson coefficient: The background noise was removed by measuring it, then subtracting it in ImageJ. The electroporated zone was then divided into three different compartments as mentioned previously. Each compartment was then analyzed using the JACoP plugin in ImageJ. An empirical threshold was used, and the Pearson’s coefficient calculated.

### All embryos which normally developed and expressing pHluo-vectors at the thoracic level were included in the analysis

Number of independent experiments, embryos, stacks and growth cones (n) are indicated in figures or legends. Analysis were done in blind for the quantification of phenotype in mouse embryos and the collapse assays. Sample size and statistical significances are represented in each figure and figure legend. For each set of data, normality was tested and Student t or Mann-Whitney tests were performed when the distribution was normal or not, respectively. Statistical tests were performed using Biosta-TGV (CNRS) and Prism 6 software.

## Acknowledgments

We thank Samir Merabet for helpful advices on the construction of fluorescent reporters, Julien Falk and Frédéric Moret for help with confocal microscopy, Camilla Lucardini, Dennis Ressnikoff, and Bruno Chapuis from the CIQLE platform of Lyon for advices on microscopy and deconvolution, Esther Stoeckli for Math1 and HoxA1 promoters constructs. STED microscopy was done in the Bordeaux Imaging Center (BIC), CNRS-INSERM-Bordeaux University, member of the national infrastructure France BioImaging supported by the French National Research Agency (ANR-10-INBS-04). We thank C. Poujol and M. Mondin (BIC) for advice, M. Sainlos for sharing the anti-GFP nanobody, and B. Tessier and S. Benquet for technical assistance. This work was conducted within the frame of the LabEx CORTEX and DEVWECAN of Université de Lyon, within the program ‘‘Investissements d’Avenir’’ (ANR-11-IDEX-0007) operated by the French National Research Agency (ANR). The study was supported by an ANR funding to VC and OT, the Association Française contre les Myopathies (AFM), the Fondation pour la Recherche Médicale (FRM) to VC, the Fondation Bettencourt-Schueller to VC, and Conseil Régional d’Aquitaine (Neurocampus funds).

## Supplemental information

**Movies S1-S4:** pHluo-PlxnA1^WT^ (S1-S2) and pHluo-PlxnA1^Y1815F^ (S3-S4) are addressed to the cell surface of commissural growth cones during the FP navigation. White arrows point the growth cones during FP navigation. FP: floor plate.

**Movies S5-S8:** FRAP sequences of commissural growth cones in spinal cord open-books. The pHluo-receptor fluorescence in an area of 15 to 20µm^2^ covering the entire growth cone surface was bleached at 80-90%. The recovery was measured over a period of 17 minutes.

**Movie S9**: 3D reconstruction with IMARIS software of axons navigating through the floor plate stained with DAPI (in blue), BEN (in white) and with mbTomato electroporated in axons (in red).

**Movie S10**: Same reconstitution as in Movie S1 but on a limited slice of spinal cord cut in the rostro-caudal axis.

**Movie S11**: The surface of a single axon from movie S2 has been reconstructed.

**Movie S12**: 3D reconstruction from a Math1-mbTomato and Hoxa1-GFP electroporation. A single Math1-mbTomato (in red) electroporated axon is navigating through the basal end-feet of a Hoxa1-GFP (in white) electroporated floor-plate cell.

**Movies S13-S16:** Sequences of time-lapse movies of chick open-books at fast time intervals (8 minutes) illustrating the navigation behaviors of PlxnA1^WT^ growth cones (S13-S14) and PlxnA1^Y1815F^ growth cones (S15-S16).

**Supplementary Figure 1:**
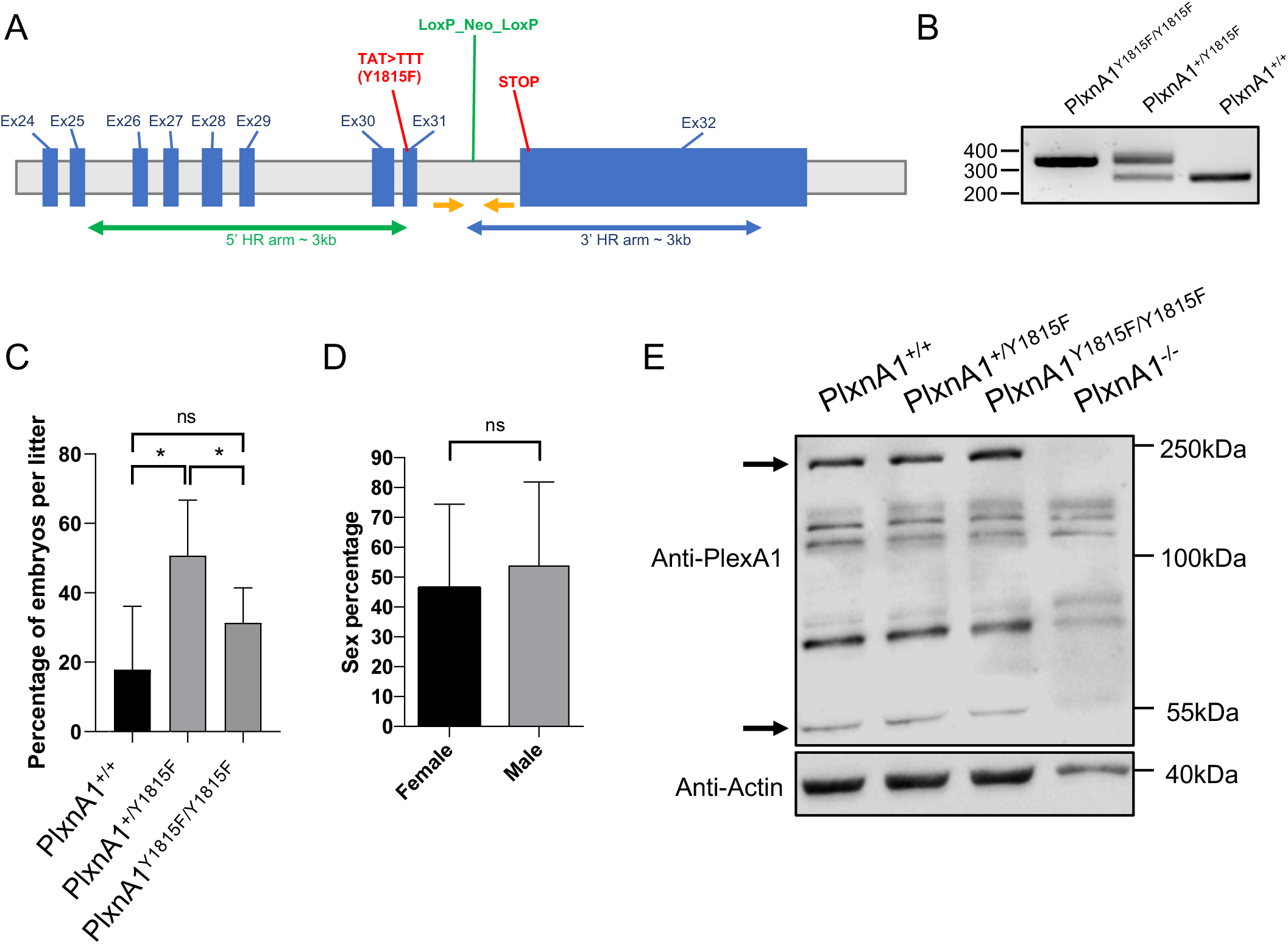
generation of PlxnA1^Y1815F^ mutant strain. (A) Schematic representation of the PlxnA1^Y1815F^ allele. The selection cassette encoding neo is inserted in the region spanning exons 25 to 32. The homolog arms are indicated in 5′ and 3′. TAT>TTT (Y1815F) mutation is inserted in intron 30-31. The genotyping primers are indicated as yellow arrows. (B) Genotyping PCR products: the genotyping primers indicated in (A) amplify a 341bp fragment from the mutated allele (PlxnA1^Y1815F/Y1815F^) and a 262bp fragment from the wild-type (PlxnA1^+/+^) allele. (C) Percentage of mice with each genotype coming from PlxnA1^Y1815F/+^ x PlxnA1^Y1815F/+^ crossing (N = 8 litters, 46 mice total). (D) Overall percentage of female and male mice (N = 41 litters, 220 mice total). Data are shown as the mean ± s.d, Student test has been applied, *: p < 0.05. (E) Representative electrophoresis of spinal cord lysates prepared from E12.5 PlxnA1^-/-^, PlxnA1^+/+^, PlxnA1^+/Y1815F^, and PlxnA1^Y1815F/Y1815F^ embryos, immunoblotted with anti-PlxnA1 and anti-actin antibodies. PlxnA1 is detected under two major forms, the integral form at 250kDa, and a short form at 55kDa. Black arrows point the 250kDa form found present at higher rate and the 55kDa form found present at lower rate in the PlxnA1^Y1815F^ condition, compared to the other genotypes.

## Notes

### Competing Interest Statement

The authors have declared no competing interest.

